# ASD gene *Bcl11a* regulates subcellular RNA localization, associative circuitry, and social behavior

**DOI:** 10.1101/2022.10.06.511159

**Authors:** Omer Durak, Ji-Yoon Kim, Dustin E. Tillman, Yasuhiro Itoh, Marit R. Wettstein, Luciano C. Greig, Thomas A. Addison, Jeffrey D. Macklis

**Author notes:** These authors contributed equally to this work (Omer Durak, Ji-Yoon Kim).

## Abstract

Precise, area-specific connectivity of interhemispheric callosal projection neurons (CPN) in the cerebral cortex is critical for sensory, associative, and behavioral functions. CPN circuitry, which connects and integrates the cerebral hemispheres via the corpus callosum, is centrally implicated in autism spectrum disorder (ASD) and intellectual disability (ID). Though transcriptional controls regulating CPN subtype and areal development are partially understood, downstream subcellular molecular machinery that implements CPN circuitry is essentially unknown. Here, we identify that the highly penetrant ASD/ID risk gene *Bcl11a/Ctip1* is critical developmentally for appropriate and precise areal targeting of superficial layer CPN (CPN_SL_) projections, and that its deletion strikingly re-routes some CPN_SL_ projections subcortically, causes dramatic disruption to subcellular CPN_SL_ growth cone (GC) molecular machinery, and disrupts social behavior and cognition in mice. CPN_SL_-specific embryonic deletion of *Bcl11a* causes loss of correct homotopic targeting in the contralateral cortex, re-routes a substantial proportion of axonal projections through the anterior commissure, and induces strikingly aberrant, but specific and precise, projections to the basolateral amygdala. Importantly, bilateral deletion of *Bcl11a* from CPN_SL_ results in abnormal social behavior and working memory. Mechanistically, we identify dysregulation of the CPN_SL_ axonal GC-localized transcriptome in *Bcl11a* nulls, due to either aberrant transcription or trafficking of individual transcripts. These molecular mis-localizations disrupt axon guidance and CPN_SL_ circuitry formation. Together, this work identifies critical functions for *Bcl11a* in CPN_SL_ axonal connectivity, development of functional associative-social circuitry, and regulation of subtype-specific subcellular molecular machinery *in vivo*, revealing novel GC-localized transcripts that regulate precise axonal targeting and circuit formation. These results elucidate development and ASD/ID disease-related circuit disruption of CPN_SL_, and the importance of understanding circuit-specific subcellular molecular machinery by neurons.

## Introduction

The mammalian cerebral cortex, which is responsible for processing sensory information, fine motor control, and higher-order cognition, has undergone dramatic expansion during evolution– largely due to increase in both number and diversity of superficial inter-hemispheric callosal projection neurons (CPN_SL_) and ipsilateral inter-areal projections neurons^1,2^. Interhemispheric commissural PN (primarily composed of CPN) are the broad population of cortical excitatory projection neurons that connect the two cerebral hemispheres by extending their primary axonal projections to homotopic regions of the contralateral hemisphere, mainly via the *corpus callosum* (CC). A small percentage of commissural neurons connect via the evolutionarily older anterior commissure (AC) tract^3,4^. CPN, through bilateral transfer and integration of cortical information, play key roles in high-level associative, integrative, cognitive, behavioral, and sensory functions, based on precise, area-specific CPN subtype connectivity and diversity. CPN evolved relatively recently in eutherians, with dramatic expansion in mammals with more complex sensorimotor integration and cognition^2,5–8^. Importantly, disruption of CPN circuitry is associated with neurodevelopmental and cognitive disorders. Individuals born with agenesis of the CC exhibit a range of cognitive and behavioral disorders, centrally including ID and ASD^9,10^. The remarkable range of cognitive and behavioral deficits associated with disruptions in CPN/CC development reflects the complex development and diversity of this population, and highlights the importance of CPN diversity and precise development in high-level cognition and associative behavior.

Recently, distinct combinatorial sets of genes have been identified both that define and control development of CPN^11^. However, little is known about molecular regulation of development of precise CPN circuitry. One recently identified molecular control over precise sensory-associative CPN connectivity is the transcription factor *Bcl11a/Ctip1*, which functions to regulate both precision of CPN areal connectivity and subtype development^12,13^. *Bcl11a* is expressed specifically by CPN (and similarly sensory processing corticothalamic PN) in all primary sensory areas of cortex, but is excluded from corticospinal/subcerebral PN. Forebrain-specific deletion of *Bcl11a* results in substantial reduction of the CC, disrupted specification of these distinct sensory-associative cortical projection neuron subtypes, and aberrant contralateral targeting^12,13^. CPN_SL_ also require *Bcl11a* for proper radial migration and localization^12,14^. Haploinsufficiency of *Bcl11a* in mice results in impaired cognition and social interaction^15^. Thus, *Bcl11a* mutant mice link gene function to CPN circuitry to behavior. Importantly, recent studies have found that patients with either microdeletion of 2p16.1, which includes *BCL11A*, or missense or loss-of-function mutations of *BCL11A* display central features of ASD, ID, and developmental delay, as well as hypoplasia of the CC and microcephaly^16–20^.

Many ASD risk genes encode proteins known to be important for synapse formation, cell-cell communication, and transcriptional regulation, suggesting that common molecular etiology might exist, at least for a subset of ASD patients^21,22^. Also, accumulating evidence indicates disrupted cortical development importantly underlying core aspects of ASD^23,24^. Further, ASD-associated risk genes are predominantly enriched in the superficial cortical layers, where most CPN reside, in human fetal cortex^25^. These results support system-level findings of disconnection in long-distance circuits in ASD^26^, with weak functional connectivity and synchronization between the cerebral hemispheres^27^. These are core CPN functions, and CPN dysgenesis is one of only a few reproducibly identified pathologies across ASD, with reduced CC connectivity relative to brain volume in many ASD patients^28^.

Thus, CPN dysgenesis with *Bcl11a* mutations is highly relevant to understanding a large subset of ASD pathology. While progress has been made identifying transcriptional controls regulating CPN subtype and areal development, downstream subcellular mechanisms and molecular machinery that implement CPN circuitry is substantially less known. Previous *in vitro* studies provide evidence that complex subcellular molecular networks are axon GC-localized, including tightly regulated local translation of select mRNA transcripts directly controlling GC pathfinding^29–31^. Recent *in vivo* work from our lab, using newly developed experimental and analytic approaches, reveals CPN_SL_ GC-specific enrichment of hundreds of transcripts and proteins compared with parent CPN somata, indicating that subcellular GC molecular machinery directly implements subtype-specific nuclear developmental programs to build circuitry, connect to appropriate targets, and select correct synaptic partners^32,33^. Molecular function of subtype- and stage-specific GCs is thus critical for understanding development, organization, maintenance, and disruption of ASD-related subtype-specific circuits *in vivo*.

In the work presented here, we identify that *Bcl11a* is critical developmentally for appropriate and precise areal targeting of CPN_SL_ associative projections– that its deletion strikingly reshapes these projections, and causes dramatic disruption to circuit-specific subcellular GC molecular machinery and social behavior and cognition in mice. Dual-population mosaic analysis of spatially and temporally identical *Bcl11a* null vs. control CPN_SL_ in the same brains identifies aberrant, non-homotopic targeting in contralateral cortex, substantial increase of projections through the AC, and remarkably specific aberrant projections to the fear and anxiety processing hub of the basolateral amygdala (BLA) in developing mouse brain, all of which persist in adult brain. Importantly, bilateral deletion of *Bcl11a* from CPN_SL_ circuitry alone results in deficient social novelty preference and working memory in both male and female mice.

Applying the recently developed approaches to isolate, purify, and quantitatively analyze subtype- and stage-specific GCs *in vivo*, we identify specific dysregulation of a set of GC-localized transcripts in *Bcl11a* null CPN_SL_ GCs, due to either aberrant transcription or trafficking of individual transcripts, including as exemplars ASD gene *PcdhαC2*, and *Mmp24*. Such disruption of the local GC transcriptome likely importantly underlies defects in axon guidance and development of CPN_SL_-specific associative circuitry, causing disease related behavior. Indeed, knockdown of *Mmp24* and *PcdhαC2* in CPN_SL_ causes disruption of precise contralateral CPN_SL_ targeting and a shift in targeting towards medial cortex. Together, this work identifies critical functions of *Bcl11a* in axonal connectivity by CPN_SL_, development of functional associative-social circuitry, and regulation of subtype-specific subcellular molecular machinery *in vivo*, thus revealing novel GC-localized transcripts as candidates regulating precise axonal targeting and circuit formation. These results elucidate development and ASD/ID-related circuit disruption of CPN_SL_, and the importance of understanding circuit-specific subcellular localization of molecular machinery by neurons.

## Results

### *Bcl11a* deletion causes aberrant and imprecise CPN_SL_ contralateral projections during development that persists into adulthood

CPN are centrally implicated in high-level associative, integrative, and cognitive behaviors through their precise and diverse area-specific interhemispheric connectivity. Building upon our previous findings that *Bcl11a* functions importantly in both cortical primary sensory area identity development and subtype specification of deep layer cortical neurons^12,13^, we asked whether *Bcl11a* also functions cell-autonomously in CPN_SL_ to regulate their axonal development and areally-precise targeting. Because CPN_SL_ axonal development and projection requires tightly regulated spatial and temporal developmental processes, to deeply investigate cell-autonomous *Bcl11a* function(s) in development of CPN_SL_ connectivity, we compared axonal projections of spatially and temporally identical and interspersed *Bcl11a* cKO (heterozygous and homozygous) and WT CPN in the same mice at multiple, critical developmental stages by applying our lab’s newly developed genetic mosaic plasmid system at three neonatal and mature ages: 1) at postnatal day 0 (P0), when CPN_SL_ begin to cross the midline; 2) at P7, during CPN homotopic axonal targeting specification, but before axonal pruning; 3) at P60, after developmental axonal pruning.

To delete *Bcl11a* specifically in CPN_SL_ unilaterally, with an intermingled set of WT CPN_SL_ within the same area of the same brain, while leaving the contralateral hemisphere intact (wildtype), we electroporated individual embryos from *Bcl11a*^flox/flox^ or *Bcl11a*^flox/+^ litter *in utero* at E14.5 with a precisely-defined cocktail of Cre-recombinase, Cre-silenced myristoylated (myr-)-EGFP (control), Cre-activated myr-tdTomato (cKO), and Cre-inducible Cre-recombinase^34^, and collected brains at P0, P7, and P60 (Fig. 1a; Supp. Fig. 1a, b). At P7, both control and *Bcl11a* null CPN_SL_ axons project to the contralateral hemisphere. Remarkably, in contrast to WT CPN_SL_ that project highly specifically to homotopic regions in S1/S2 with distinct bimodality, interspersed *Bcl11a* null CPN_SL_ project promiscuously across the contralateral cortex without focus or specificity, and substantially increase far lateral projections, peaking in highly inappropriate insular cortex (Fig. 1b, c). To investigate whether early postnatal axonal targeting disruption is potentially corrected by axonal pruning later in development, we further asked whether the loss of areal precision of CPN_SL_ axonal projections at P7 persists and causes long-lasting changes in cortical circuitry in adult cortex. We analyzed CPN_SL_ axonal targeting across the medio-lateral extent of contralateral cortex in P60 brains.

**Figure 1:**
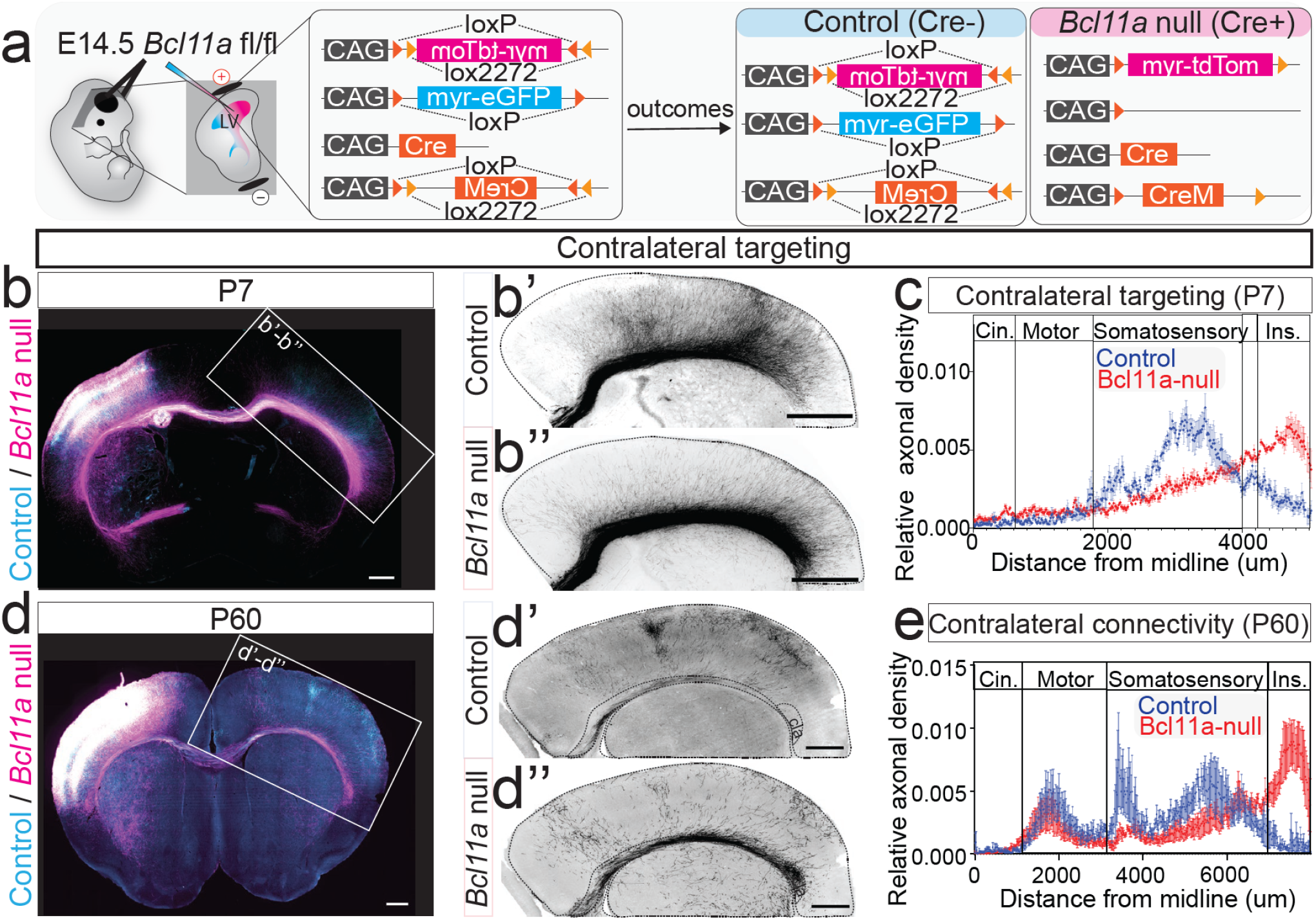
*Bcl11a* deletion from superficial layer callosal projections neurons (CPN_SL)_ disrupts precise associative connectivity. **(a)** Schematic for dual-population mosaic targeting of CPN_SL_ *in vivo*^34^. LV = lateral ventricle. **(b)** Fluorescence images of P7 *Bcl11a*^flox/flox^ mouse brain electroporated *in utero* at E14.5 with a precisely-defined cocktail of Cre-recombinase, Cre-silenced myr-eGFP expression construct (control; cyan), Cre-activated myr-tdTomato (*Bcl11a* null; magenta), and Cre-inducible Cre-recombinase. High magnification greyscale images of contralateral cortex for control (b’) and *Bcl11a* null (b”). **(c)** Quantification of contralateral axonal targeting at P7 reveals disruption of contralateral homotopic targeting of *Bcl11a* null CPN_SL_ axons compared to control (Control, n=5; *Bcl11a* null, n=5). **(d)** Fluorescence images of P60 *Bcl11a*^flox/flox^ mouse brain electroporated *in utero* at E14.5 (control, cyan; *Bcl11a* null, magenta). High magnification greyscale images of contralateral cortex for control (d’) and *Bcl11a* null (d”). cla = claustrum. **(e)** Quantification of contralateral axonal targeting at P60 reveals that loss of areal precision of associative projections observed during early cortical development persists into adulthood (Control, n=4; *Bcl11a* null, n=4). All sections were immunolabeled for GFP (control; cyan) and RFP (*Bcl11a* null; magenta). Scale bars are 1 mm.

As with P7 CPN_SL_ projections, there is similar loss of specificity or clustering of axonal projections in somatosensory cortex, normally seen in wt cortex, following *Bcl11a* deletion, as well as striking increase in aberrant targeting of insular cortex (Fig. 1d, e). These results suggest that *Bcl11a* deletion causes CPN_SL_ to lose areal precision of associative projections by CPN_SL_-autonomous, aberrant axonal targeting during early cortical development, which importantly persists into adulthood.

Importantly, this disruption of contralateral axonal targeting is not due to stalled axonal elongation or defective midline crossing, since *Bcl11a* null CPN_SL_ axons project across the midline to contralateral cortex at P0, when WT somatosensory CPN_SL_ have only begun to cross the mideline through the CC (Supp. Fig. 1f). Notably, the ratio of axons crossing the midline is higher for null CPN_SL_ compared to WT even after normalizing to the pre-midline axonal density (Supp. Fig. 1g). This indicates that *Bcl11a* deletion does not disrupt axonal elongation and midline crossing processes of CPN_SL_, but might even increase elongation speed, potentially via loss of regulation for target precision.

Extending our lab’s previous results revealing that forebrain-specific deletion of *Bcl11a* causes enlargement of the AC due to an increased number of cortical neurons projecting through this axonal tract^12^, we observed that many *Bcl11a* null CPN_SL_ in somatosensory cortex also aberrantly project CPN_SL_-autonomously to the contralateral hemisphere through the AC, as compared to wt (Fig. 2a, b). In addition to predominantly projecting through the CC, a small minority of interhemispheric connections project through the evolutionarily older AC, connecting homotopic areas in piriform and entorhinal cortices, as well as amygdaloid regions^35^. Our results indicate that *Bcl11a* functions cell-autonomously to control the interhemispheric axonal projection trajectory of CPN_SL_ in developing mouse cortex, causing them to project via the evolutionarily newer CC. Further, and quite strikingly, in addition to the abnormal contralateral axonal projections via the AC and essentially total disruption of homotopic targeting in the absence of *Bcl11a* function, *Bcl11a* null CPN_SL_ form highly specific bilateral projections to the basolateral amygdala (BLA), which are essentially absent in WT brains (Fig. 2c, d).

**Figure 2:**
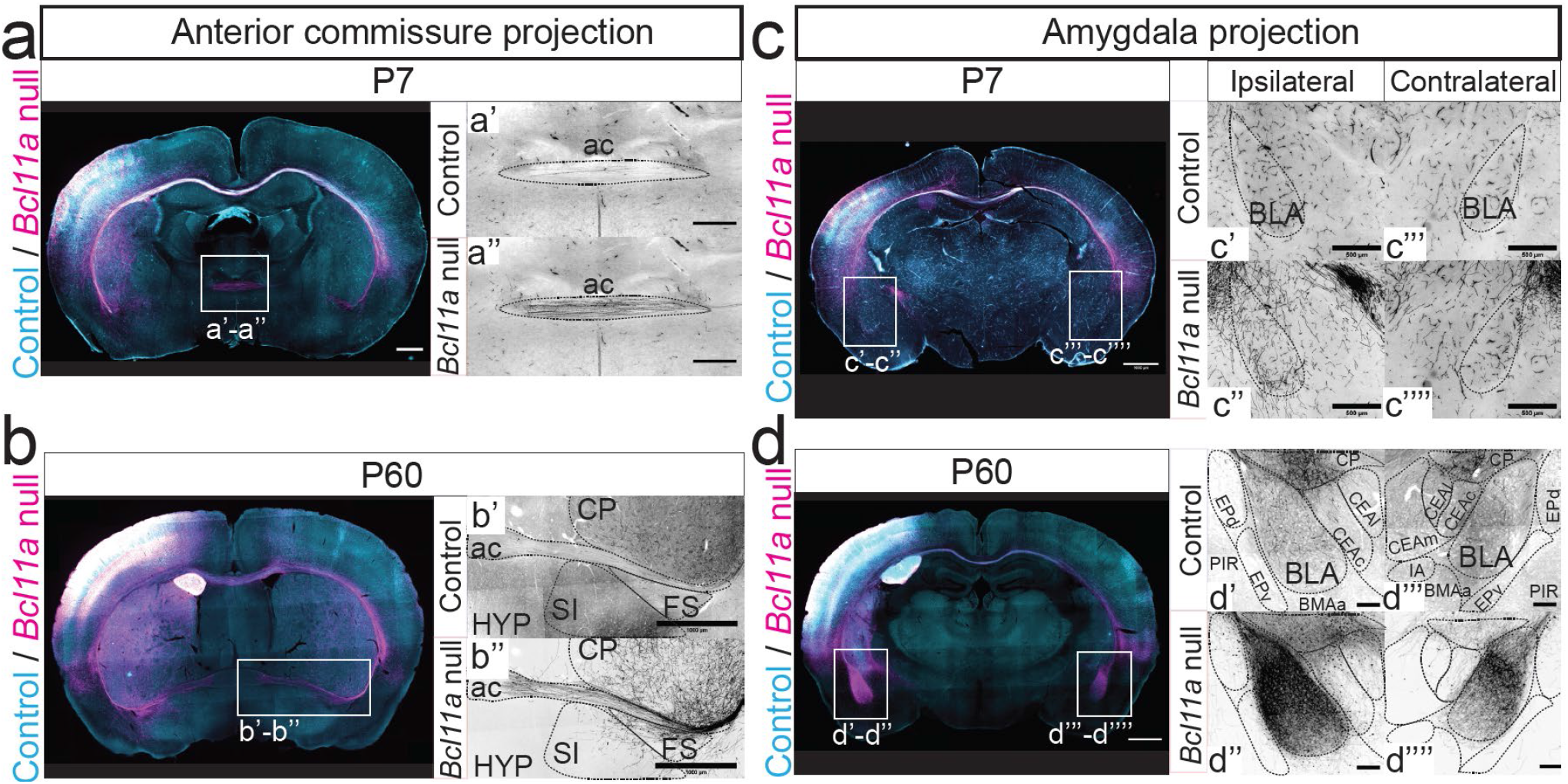
*Bcl11a* deletion from CPN_SL_ aberrantly increases projections through the AC and causes aberrant *de novo* projections to BLA. **(a)** Fluorescence images of P7 brain with mosaic population of *Bcl11a* null (magenta) and control (cyan) reveal increased aberrant CPN_SL_ axonal projections through the AC upon *Bcl11a* deletion. High magnification greyscale images of AC for control (a’) and *Bcl11a* null (a”). **(b)** Fluorescence images of P60 brain reveal that increased aberrant axonal projections through AC upon *Bcl11a* deletion persist into adulthood. High magnification greyscale images of AC for control (b’) and *Bcl11a* null (b”). **(c)** Fluorescence images of P7 brain reveal that *Bcl11a* null CPN_SL_ axons already start to enter the ipsilateral BLA at P7, whereas very few control axons do. High magnification greyscale images of BLA for control (c’, ipsilateral; c”‘, contralateral) and *Bcl11a* null (c”, ipsilateral; c”“, contralateral). **(d)** Fluorescence images of P60 brain reveal that *Bcl11a* null CPN_SL_ form highly specific bilateral projections to BLA compared to control. High magnification greyscale images of BLA for control (d’, ipsilateral; d”‘, contralateral) and *Bcl11a* null (d”, ipsilateral; d”“, contralateral). All sections were immunolabeled for GFP (control; cyan) and RFP (*Bcl11a* null; magenta). Scale bars are 1 mm, except 500µm for high magnification greyscale images.

To eliminate any potential confounding effects due to differential levels of green vs. red fluorescent labeling, we investigated WT vs *Bcl11a* null CPN_SL_ projections at P7 by inverting fluorescence labeling between the two conditions (i.e. control with tdTomato, and cKO with EGFP), and observed the same circuit defects using this inverted fluorescence color system (Supp. Fig. 2). Finally, heterozygous deletion of *Bcl11a* from CPN_SL_ does not equivalently disrupt contralateral cortex targeting, but does cause increased projections through the AC at P7, which recede by P60 (Supp. Fig. 3).

Together, our results indicate that Bcl11a is critical for proper integrative association between the hemispheres, by controlling precise targeting and refinement of connectivity between homotopic locations in the cerebral hemispheres. Further, homozygous deletion of *Bcl11a* in CPN_SL_ alone, with an otherwise wt, normal brain, causes aberrant, but highly specific and precise, projections to multiple pathological regions, including BLA and via the anterior commissure, circuit-level disruptions implicated specifically in cognitive and social behaviors.

### CPN_SL_-specific deletion of *Bcl11a* during development impairs social novelty preference and working memory in adult mice

Our results at the level of connectivity indicate that *Bcl11a* is critical for proper integrative association between the hemispheres and to prevent incorrect projections to multiple regions involved in human social behavioral and cognitive pathology. Importantly, the aberrant projections by *Bcl11a* null CPN_SL_ observed in early postnatal development persist into adulthood, implicating circuit-level interference in adult behavior. Strong correlation between ASD-related cognitive and behavioral deficits and CPN/CC dysgenesis^28,36–39^ highlight the importance of precise CPN circuitry development in social behavior and cognition. Therefore, we hypothesized that the observed abnormalities of *Bcl11a* null CPN_SL_ projections might underlie ASD-associated behavioral features in these mice.

To elucidate whether absence of *Bcl11a* function in CPN_SL_ alone results in ASD-related aberrant social behavior and cognition, we bilaterally *in utero* electroporated individual embryos from *Bcl11a*^*fl/fl*^ litters at E14.5 with either a control myr-EGFP reporter or a myr-tdTomato reporter and Cre, targeting CPN_SL_. Bilateral targeting and successful labeling of CPN_SL_ was confirmed following the completion of the behavioral assays (Supp. Fig. 4). In the rostro-caudal axis, we observed robust labeling in somatosensory and motor cortices, with minimal coverage in visual cortex, confirming successful bilateral CPN_SL_- and area-specific *Bcl11a* deletion for behavioral interrogation.

We completed an extensive set of behavioral assays widely employed to investigate what are thought to be ASD-related and ID-relevant behavioral assays, covering social interaction, exploratory behavior, learning and memory/cognition, and stereotyped behavior. We conducted behavioral examination with both males and females (Fig. 3a). First, to test for potential confounding effects that could theoretically arise due to abnormal gross locomotor activity and/or exploration habits, we employed an open field test to observe freely moving mice in an empty arena. We observed similar basic locomotor activity between *Bcl11a* CPN_SL_ cKO and control mice, measuring several variables: total distance travelled, velocity, total time in motion, and time spent in the center of the chamber (Supp. Fig. 5a-d). Further, we examined the conflict between exploration and risk avoidance using both a light-dark box and an elevated plus maze ^15,40^. Both male and female *Bcl11a* CPN_SL_ cKO mice were unchanged compared to controls; they displayed no significant differences in the number of entries to, or the amount of time spent, in the light chamber or the open arms (Supp. Fig. 5e-h). These data indicate that *Bcl11a* deletion from bilateral CPN_SL_ in somatosensory and motor areas does not affect basic locomotory activity or exploration, providing a rigorously unconfounded foundation for the more specific behavioral experiments to follow.

**Figure 3:**
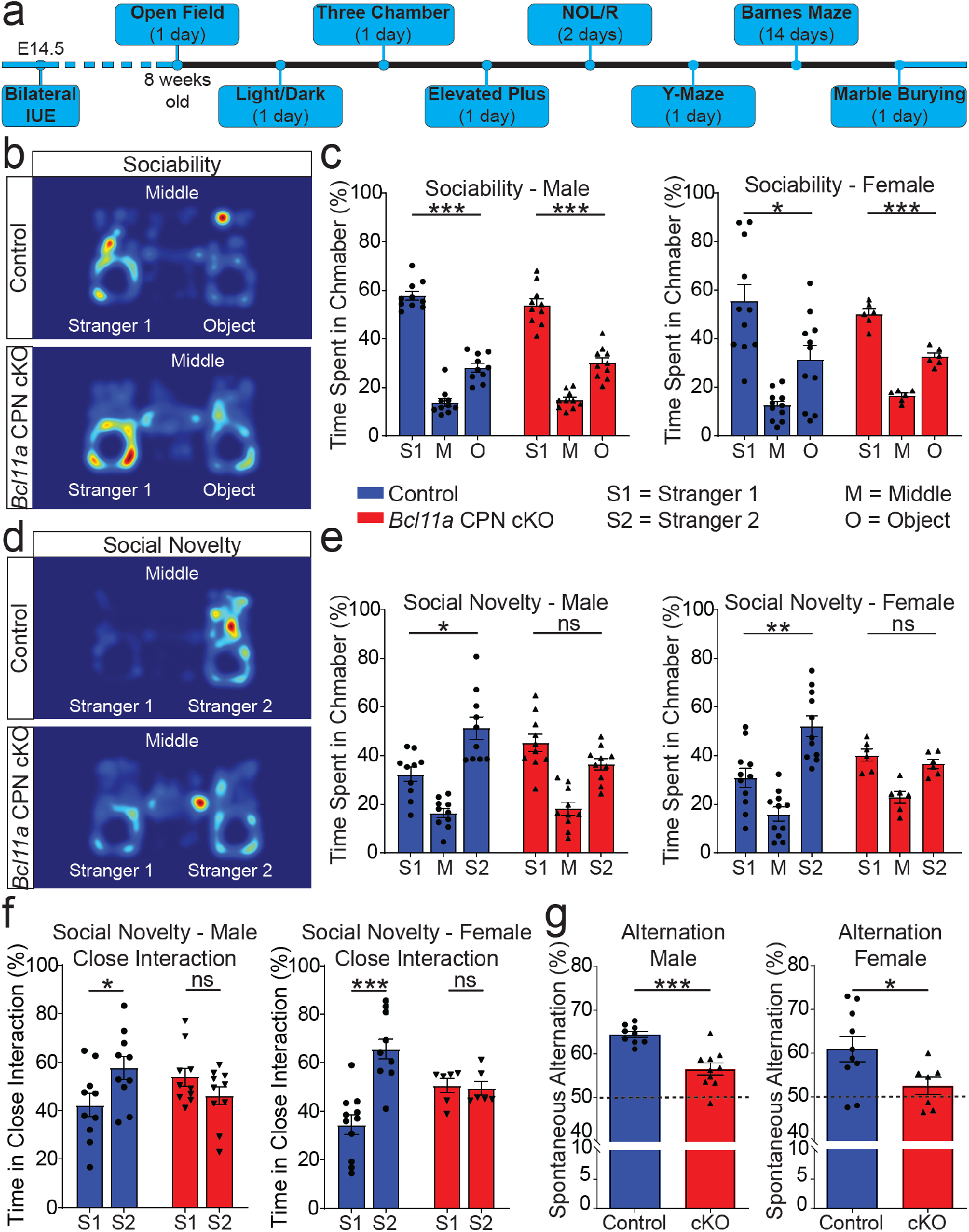
CPN_SL_-specific deletion of *Bcl11a* selectively impairs social novelty preference and working memory. **(a)** Timeline and durations of eight distinct behavioral experiments conducted after bilateral *in utero* electroporation (IUE) of Cre recombinase at E14.5 to delete *Bcl11a* from CPN_SL_. **(b)** Representative heat map analyses of sociability testing of control and *Bcl11a* CPN_SL_ cKO mice. **(c)** Quantification of sociability reveals no significant difference between *Bcl11a* CPN_SL_ cKO and control mice (both males and females; one-way ANOVA followed by Bonferroni multiple comparison test; control male, n=10; cKO male, n=10; control female, n=11; cKO female, n=6). **(d)** Representative heat map analyses of social novelty preference testing of control and *Bcl11a* CPN_SL_ cKO mice. **(e)** Quantification of social novelty preference for both males (left) and females (right) reveals that *Bcl11a* CPN_SL_ cKO mice lack preference for the new stranger (Stranger 2) compared to control mice (one-way ANOVA followed by Bonferroni multiple comparison test; control male, n=10; cKO male, n=10; control female, n=11; cKO female, n=6). **(f)** Both male and female *Bcl11a* CPN_SL_ cKO mice display no preference for the new stranger (Stranger 2) in close interaction analysis of social novelty preference (two-tailed Student’s t test; control male, n=10; cKO male, n=10; control female, n=11; cKO female, n=6). **(g)** Quantification of spontaneous alternation in Y-maze test reveals that both male (left) and female (right) *Bcl11a* CPN_SL_ cKO mice make significantly fewer alternations compared to controls (two-tailed Student’s t test; control male, n=9; cKO male, n=10; control female, n=10; cKO female, n=7). *, p-value<0.05; **, p-value<0.01; ***, p-value<0.001; ns, nonsignificant. n, number of distinct mice used. Results are represented as mean±s.e.m.

Abnormal social interactions constitute the core behavioral deficits in ASD, along with cognitive deficits and repetitive behavior in many ASD subtypes, including those with *Bcl11a* deletion/mutations^15,40^. To investigate whether the absence of *Bcl11a* function in CPN_SL_ alone modifies social behavior in mice, we employed a three-chamber social arena assay to examine the sociability and social novelty. During the habituation period, both *Bcl11a* CPN_SL_ cKO and control mice explored the lateral chambers equally (Supp. Fig. 6a), further reinforcing no overall change in exploratory behavior. Further, both groups displayed equivalent sociability– preference for time spent in the lateral chamber with the stranger mouse vs. the inanimate object, as well as in close interaction (Fig. 3b-c, and Supp. Fig. 6b). Remarkably, when the inanimate object was replaced with a novel stranger mouse, both male and female *Bcl11a* CPN_SL_ cKO mice showed no preference for the new stranger over the first stranger, whereas control mice spent significantly more time in the chamber with the novel mouse, and in close interaction with the cage (Fig. 3d-f). These results indicate strongly that *Bcl11a* function in CPN_SL_ alone, thus correct CPN_SL_ circuit development, are critical for intact social novelty preference, whereas *Bcl11a* function in CPN_SL_ circuitry is dispensable for general sociability.

In addition to abnormal social behavior, impaired cognition of variable degree is a hallmark of many subsets of ASD, and *Bcl11a* deletions/mutations cause intellectual disability in humans. CPN circuitry performs high-level associative, integrative, and cognitive function via precise and diverse area-specific connections. The prior experiments reveal that CPN circuitry is substantially disrupted in the absence of *Bcl11a* function. Therefore, we hypothesized that specific abnormalities of learning, memory, and/or cognition would result from the area-specific *Bcl11a* deletion from CPN_SL_, rather than wide decline in cognition. To investigate cognition both widely and more specifically, we investigated working memory, spatial learning and memory, and recognition memory via a Y-maze, a Barnes maze, and a novel object recognition test, respectively. As predicted, mice with bilateral *Bcl11a* CPN_SL_ deletion exhibit specific disruption of working memory, but no change in spatial learning and memory, or novel object recognition (Supp. Fig. 7). In Y-maze spontaneous alternation test, all groups had equivalent exploration, with a similar number of total number of entries to the arms. However, control mice performed significantly better than chance (50% spontaneous alternation), as expected, whereas mice with bilateral *Bcl11a* CPN_SL_ deletion performed significantly worse compared to control group (Fig. 3g).

These results reveal that bilateral deletion of *Bcl11a* from sensorimotor area CPN_SL_ alone causes deficient social novelty preference and/or discrimination, along with working memory deficits, without major changes in other classical cognitive and/or ASD-associated behaviors. Thus, *Bcl11a* function in CPN_SL_ during circuit establishment, and proper CPN_SL_ circuitry by extension, is important specifically for social novelty preference behavior and working memory, strongly linking CPN_SL_ circuitry to these ASD-associated aberrant behaviors.

### Bcl11a controls both correct gene expression and subcellular localization of axon GC molecular regulators in developing CPN

Our results from unilateral deletion of *Bcl11a* from CPN_SL_ identified that CPN lose areal precision of associative projections by cell-autonomous, aberrant axonal targeting, rather than by loss of target area identity. These results strongly indicate that the causal abnormalities of molecular machinery responsible for axonal pathfinding and target engagement are highly likely localized to *Bcl11a* null CPN_SL_ GCs. We hypothesized that, in addition to previously identified *Bcl11a* functions in transcriptional regulation in CPN_SL_ somata^13^, *Bcl11a* might also regulate CPN_SL_ subcellular axonal GC molecular machinery by mechanisms separate from transcriptional regulation. Together, the prior data suggest that both transcriptional and potentially distinct, non-transcriptional, such as subcellular trafficking, *Bcl11a* functions together critically regulate associative cortical circuit development, maintenance, and behavior.

We directly investigated whether there might be dysregulation of subcellular GC vs. soma RNA localization in *Bcl11a* null CPN_SL_ using recently developed experimental and analytic approaches for quantitative “subcellular RNA mapping” of quite distinct GC vs. soma transcriptomes^32,33^. CPN_SL_ GCs contain up to hundreds of unique RNAs (and proteins) enriched significantly and specifically compared to their own parent somata. We simultaneously isolated *Bcl11a* null, heterozygous, and WT CPN_SL_ GCs and their parent somata^32^. In brief, to purify subtype-, stage-, and area-specific sensorimotor CPN_SL_ GCs and their parent somata from the same cortices, individual embryos from *Bcl11a*^flox/flox^ or *Bcl11a*^flox/wt^ litters were electroporated *in utero* at E14.5 with myr-tdTomato, nuclear-GFP, and Cre constructs for *Bcl11a* null or heterozygous samples, or myr-tdTomato and nuclear-GFP for wt samples. Fluorescent CPN_SL_ GCs from each condition were purified from contralateral cortices at P3 by established methods^32,33^ for subcellular fractionation and fluorescent small particle sorting (FSPS; Fig. 4a). Matching fluorescent parent somata were isolated from ipsilateral cortices by neuronal FACS^32,33^ (Fig. 4a). To confirm that the subcellular fractionation enriched for GCs prior to their purification, we compared GAP43 (GC marker) and MAP2 (somato-dendritic marker) protein abundance in the GC fraction (GCF) vs. “post-nuclear homogenate” (input; PNH) via western blot. As previously reported^32,33^, there was substantial enrichment of GAP43 and de-enrichment of MAP2 in the GCF (Supp. Fig. 8a). Further, to confirm that isolated GCs rapidly re-sealed and were intact, surrounding their RNA and protein contents, we performed RNase and protease protection assays. Extracted RNA and GAP43 protein was protected in the detergent-only or enzyme-only conditions compared to the detergent+enzyme condition, indicating that isolated GCs have sealed and intact membranes (Supp. Fig. 8b). Following purification of GCs and somata, RNA was extracted and sequenced to investigate the subcellular transcriptomes. All transcriptomes clustered by condition (Supp. Fig. 9a, b), and were further assessed by rigorous quality control (see Materials and Methods).

**Figure 4:**
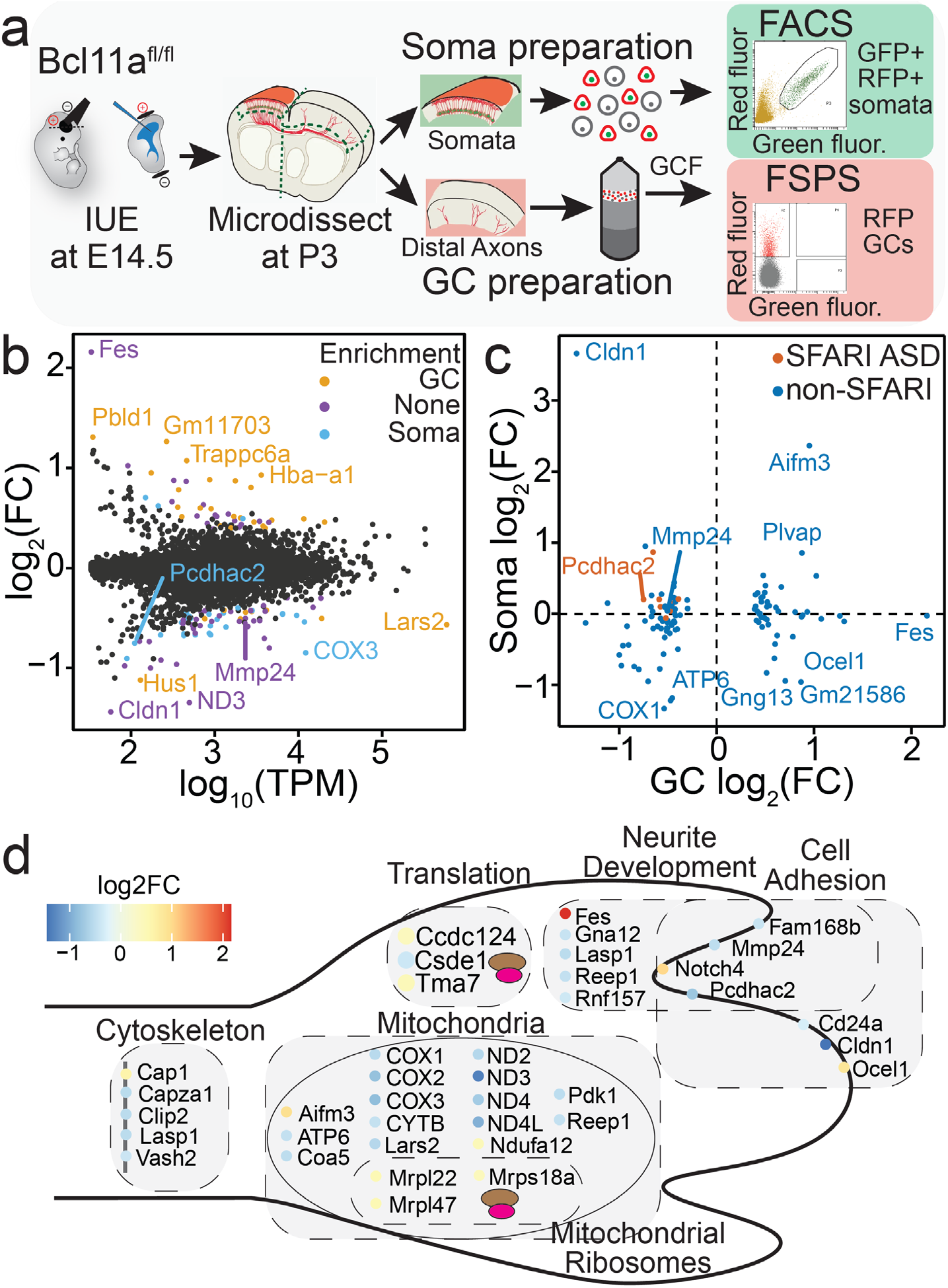
*Bcl11a* controls expression and subcellular localization of CPN_SL_ GC mRNAs. **(a)** Schematic for purification of CPN_SL_ GCs and their parent somata. **(b)** MA-plot showing changes in gene abundance in CPN_SL_ GCs upon deletion of *Bcl11a*. Differentially abundant genes (DAGs) are colored by their subcellular enrichment in WT CPN_SL_ GCs, somata, or neither (Supp. Fig. 9c) - GC-enriched DAGs are orange, soma-enriched are blue, and non-enriched are purple. **(c)** Quadrant plot comparing changes in abundance of GC DAGs in CPN_SL_ somata and GCs upon deletion of *Bcl11a* specifically from CPN_SL_. DAGs are colored by their SFARI classification – SFARI-listed ASD risk genes are red, and non-listed genes are blue. **(d)** Schematic for functional categorization of GC DAGs. DAGs that increase in CPN_SL_ GCs upon *Bcl11a* deletion have warmer colors, and DAGs that decrease in CPN_SL_ GCs upon *Bcl11a* deletion have cooler colors.

A focused set of 113 protein-coding genes in *Bcl11a* null GC samples were identified as “differentially abundant genes” (DAGs; agnostic regarding mechanism, whether via expression or trafficking, e.g.) compared to control GC samples (q-value < 0.05; Fig. 4b). To determine whether the DAGs are enriched for genes normally more abundant in GCs compared to somata, we conducted enrichment analysis between WT soma and WT GC samples to categorize genes as soma-enriched, GC-enriched, or unenriched (Supp. Fig. 9c). Quite strikingly, this quantitative subcellular analysis reveals that many normally soma-enriched genes tend to become less abundant in GCs after *Bcl11a* deletion, whereas normally GC-enriched genes become more abundant in null GC samples compared to control GC samples (Fig. 4b). These results indicate that molecular mechanisms regulating targeted mRNA trafficking to, and enrichment in, axonal GCs become more stringent, resulting in aberrant subcellular localizations, distributions, and “mapping” of many specific mRNA transcripts (Fig. 4b).

To investigate this finding further, we determined expression levels of DAGs between *Bcl11a* null and WT CPN_SL_ somata to distinguish individual transcripts whose trafficking to GCs appears to be: 1) specifically reduced (i.e., increased expression in somata, reduced abundance in GCs); 2) actively increased (i.e. reduced expression in somata, increased abundance in GCs); and/or 3) secondarily dysregulated in GCs due to differential somatic expression (Fig. 4c). Remarkably, approximately half of the DAGs are dysregulated in null CPN_SL_ GCs in the opposite direction of their somatic expression changes, strongly implying that their trafficking to, and abundance in, GCs is controlled actively and with specificity, rather than by, e.g., indiscriminate mechanisms for all GC-localized transcripts. Together, these results indicate that *Bcl11a* is necessary for correct trafficking of many specific mRNAs to sensorimotor CPN_SL_ axonal GCs during development, disruption of which likely causes, in aggregate, the striking *Bcl11a* null CPN_SL_ circuit dysgenesis and behavioral dysfunction. We thus investigated whether DAGs tend to encode for genes known to be important for and/or associated with axonal development. Indeed, DAGs include groups encoding proteins associated with neurite development, cell-cell adhesion, cytoskeleton/cytoskeletal regulation, mitochondrially-encoded/mitochondrial function, mitochondria ribosomal proteins, translation and protein processing/trafficking/transport/ubiquitination (Fig. 4d).

Quite notably, the data reveal enrichment of genes strongly associated with ASD risk by the Simons Foundation Autism Research Initiative (SFARI) among the transcripts with reduced GC abundance in *Bcl11a* null GCs (*PcdhαC2, Cacnb1, Clip2, Csde1, Elovl2, Elp2, Nfx1*, and *Snd1;* Fig. 4c). This striking result of a decrease in GC abundance for all ASD risk DAGs after *Bcl11a* deletion (*q-value* = 0.029, hypergeometric test) strongly supports a new mechanistic interpretation that, in addition to disproportionate transcriptional disruption of other ASD risk genes by ASD-associated transcriptional regulators reported by others, some or many ASD risk genes might likely also or instead cause pathology by disrupting subcellular localization and spatially-appropriate abundances of transcripts of other ASD risk genes, which would be locally translated into functional proteins for proper circuit development, maintenance, and function. Of further interest, seven DAGs are associated with ID (*ATP6, COX1-3, Elp2, Ndufa12*, and *Pigt;* Supp. Fig. 9d).^41^ Notably, six of these ID risk DAGs decrease in GC abundance after *Bcl11a* deletion, and five are associated with mitochondria (*ATP6, COX1-3*, and *Ndufa12*). Further, mitochondrial GO terms are also overrepresented among DAGs, suggesting that this key regulator of axon development might be disrupted upon deletion of *Bcl11a*, leading to aberrant projections (Supp. Fig. 9e).

Importantly, these RNA mis-localizations are quite specific; as predicted from the lack of gross abnormalities of axonal development, *Bcl11a* null sensorimotor CPN_SL_ GCs display preservation of core axon GC molecular machinery^32,42^ (Supp. Fig. 9f). Together, these results indicate that loss of *Bcl11a* function causes quite specific and impactful dysregulation of subcellular localization for a set of transcripts likely to function together locally in GCs in the multiple stages of precise CPN_SL_ axonal connectivity, targeting, synapse formation, maintenance, and function, ultimately causing altered behavior.

### Mmp24 and PcdhαC2 are necessary for precise CPN contralateral targeting

The results mapping RNA transcript localization of DAGs in GC vs. soma samples reveal that transcript abundances in GCs can be regulated independent of transcriptional changes in their parent somata. This strongly indicates that soma-centric analyses can miss detection and/or misinterpret changes of subcellularly dysregulated transcripts whose expression either does not change or changes in the opposite direction in the somata. We investigated two exemplar candidates among the DAGs with abundance in GCs displaying transcription-independent regulation, and for which dysregulation in GCs would be predicted to disrupt CPN axonal targeting. Among the many genes in this category, we selected *Mmp24* and *PcdhαC2* (the set also includes, e.g. *Claudin-1, Capza1, Notch4*, and *Ocel1*) whose abundances in null GCs are reduced without significant changes in somata (Fig. 4c). These candidates have substantial evidence in the literature supporting general function in neurite development, motivating investigation of their potential specific functions in CPN_SL_ axon targeting.

*Mmp24* (*MT5-MMP)* encodes a member of the matrix metalloproteinase family. This family of proteins functions in cleavage of most extracellular matrix (ECM) components. Substrates of Mmp24 include laminin-1, CSPG, N-cadherin, and fibronectin. Unlike most ECM secreted MMPs, Mmp24 localizes to cell surfaces, belonging to the transmembrane-type MT-MMPs^43^. Mmp24 is expressed mainly by neurons and oligodendrocytes, and it has been shown to localize to GCs in cultured DRG and hippocampal neurons, regulating axonal growth^43^. This suggests that Mmp24 likely functions in CPN_SL_ axon outgrowth, guidance, and homotopic targeting *in vivo* by regulating cleavage of receptors, ligands, and ECM components.

To investigate whether Mmp24 is important for precise CPN_SL_ contralateral targeting, we knocked down *Mmp24* expression in CPN_SL_ using the CRISPR-CasRx system via *in utero* electroporation at E14.5 and investigated targeting at P7. We achieved approximately 70% knockdown of *Mmp24* mRNA in sorted P3 CPN_SL_, confirming efficient targeting (Supp. Fig. 10a). Both control and *Mmp24* knockdown CPN_SL_ axons project to the contralateral hemisphere at P7, suggesting no broad disruption of axonal outgrowth. Strikingly, however, knockdown of *Mmp24* causes disruption of the distinctly clustered innervation present for control CPN_SL_, with increased aberrant medial and lateral targeting compared to control CPN_SL_ (Fig. 5a-c). An independent approach using small hairpin RNA (shRNA) targeting *Mmp24* resulted in increased medial targeting compared to control, and reduction in the lateral innervation cluster corresponding to the somatosensory cortex S1 upper lip region (Supp. Fig. 11a-c), despite somewhat less efficient *Mmp24* expression knockdown compared to the CRISPR-CasRx approach (∼55% vs ∼70%, respectively; Supp. Fig. 10b). These results reveal that *Mmp24* expression by CPN_SL_ GCs is critical cell-autonomously for precise contralateral areal targeting of CPN_SL_.

**Figure 5:**
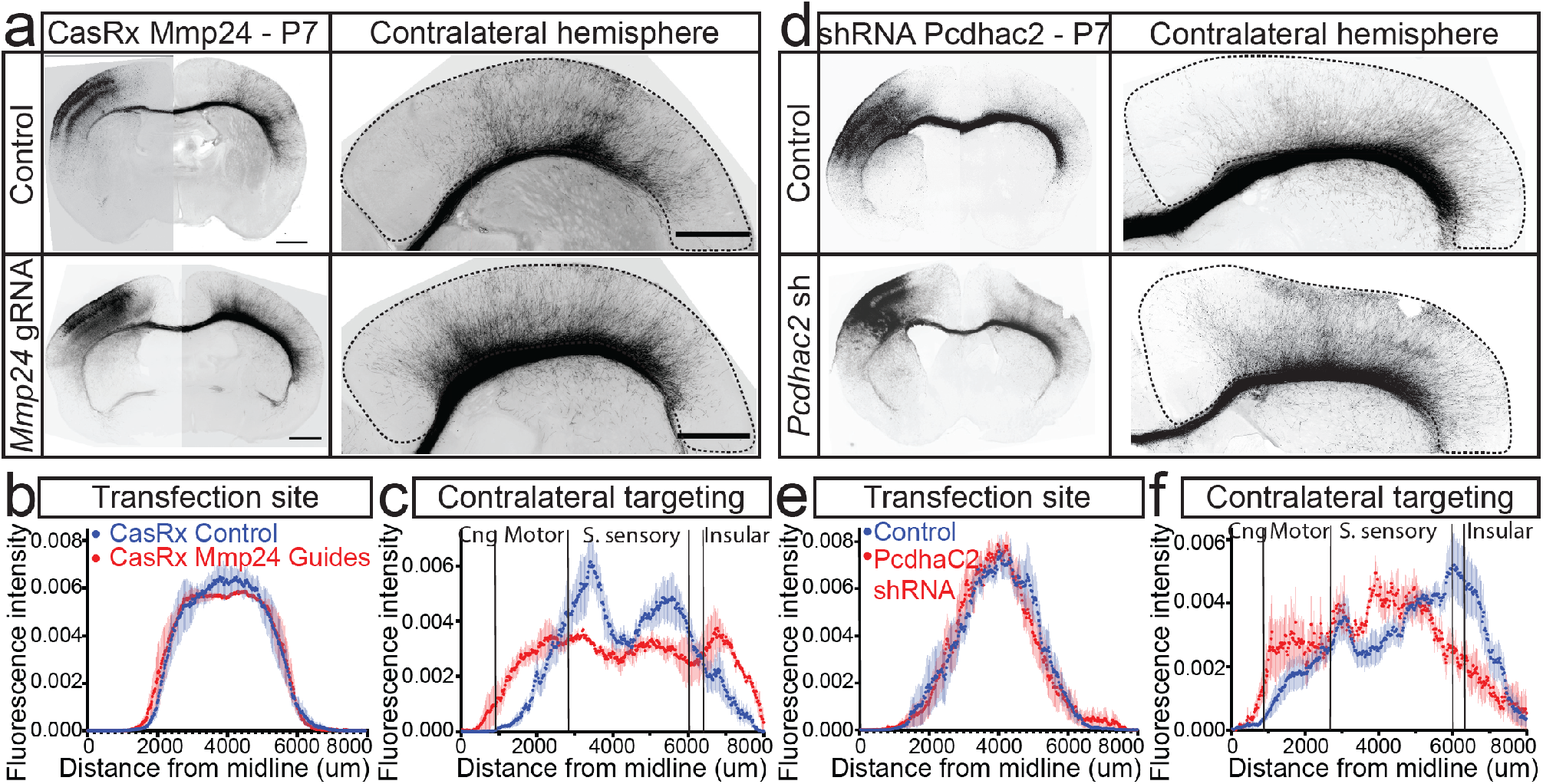
Mmp24 and PcdhαC2, dysregulated in *Bcl11a* null CPN_SL_ GCs, are necessary for precise CPN contralateral targeting. **(a)** Greyscale fluorescence images of P7 wildtype mouse brains electroporated *in utero* at E14.5 with either a non-targeting (control) or *Mmp24*-directed guide RNA, and a myr-tdTomato expression construct. Right panels are high magnification images of contralateral cortex for control (upper) and *Mmp24* gRNA (lower). **(b)** Spatial distributions of ipsilateral targeting reveal highly equivalent mediolateral transfection coverage between control and *Mmp24* gRNA CPN_SL_ somata via *in utero* electroporation (control, n=4; *Mmp24* gRNA, n=3). **(c)** Quantification of contralateral axonal targeting at P7 reveals substantial disruption of homotopic targeting of *Mmp24 gRNA* CPN_SL_ axons compared to control, with more widely and aberrantly more even distribution of targeting compared to normally more restricted, bimodal distribution (control, n=4; *Mmp24* gRNA, n=3). **(d)** Greyscale fluorescence images of P7 wildtype mouse brains electroporated with either a non-targeting (control) or *PcdhαC2*-directed shRNA (*PcdhαC2* sh), and a myr-tdTomato expression construct. High magnification images of contralateral cortex for control (upper) and *PcdhαC2* shRNA (lower). **(e)** Spatial distributions of ipsilateral targeting reveal highly equivalent mediolateral transfection coverage between control and *PcdhαC2* shRNA CPN_SL_ somata (control, n=7; *PcdhαC2* shRNA, n=7). **(f)** Quantification of contralateral axonal targeting at P7 shows substantial disruption of homotopic axonal targeting of *PcdhαC2* shRNA CPN_SL_ axons compared to control, with medial shift and spatial compaction of the normal multimodal distribution (control, n=7; *PcdhαC2* shRNA, n=7). “n” refers to number of distinct mice used. All sections were immunolabeled for RFP. Scale bars: 1 mm.

PcdhαC2 is a member of the cell membrane-localized clustered protocadherin family. These are neural cadherin-like cell adhesion proteins, expressed highly by neurons, and are known to localize to axons and synapses. Pcdhs are known to function as molecular “barcodes” for self-recognition/avoidance and tiling^44^, implying that they play important roles in establishment of subtype-specific circuits. In particular, it has been shown that *PcdhαC2* is required for axonal tiling and assembly of serotonergic circuitry^45^. With this background, we hypothesized that *PcdhαC2* might also function in precision and spatial organization of homotopic contralateral targeting by CPN_SL_ *in vivo*.

Similar to the *Mmp24* knockdown approach, we targeted *PcdhαC2* expression in developing CPN_SL_ with an shRNA that reduced *PcdhαC2* mRNA by approximately 60% in sorted P3 CPN_SL_ (Supp. Fig. 10c). Knockdown of *PcdhαC2* in CPN_SL_ causes disruption of the lateral innervation cluster and substantial increase in medial targeting in contralateral cortex compared to control shRNA (Fig. 5d-f). Knockdown of *PcdhαC2* using the CRISPR-CasRx system, less efficiently reduced *PcdhαC2* mRNA levels by ∼40% (Supp. Fig. 10d), also caused specific reduction of the lateral innervation cluster (Supp. Fig. 11d-f). Together, the results from knockdown of the exemplars *Mmp24* and *PcdhαC2* in CPN_SL_ *in vivo* both identify specific function of each of these regulators in precise areal targeting of homotopic CPN circuitry, and indicate more broadly that investigation of subcellular GC molecular machinery enables discovery of actively GC-localized transcripts that function as combinatorial subcellular molecular controls regulating precise axonal targeting and circuit formation.

## Discussion

Precise long-range and local connectivity, with area and target neuron-specific circuitry, is critical for information processing and multimodal integration in the neocortex, and in the nervous system more broadly. As an example, subtle disruption of CPN circuitry can underlie striking behavior and disability in human disorders, while leaving gross structures intact (e.g. ASD, ID, schizophrenia^44–47^), highlighting the importance of precise associative functions in computing and manipulating higher-order information. Understanding molecular mechanisms that regulate precision of circuit development and selective synapse formation by GCs, of exemplar CPN_SL_ connectivity in particular, is central to elucidating complex CNS function and dysfunction. While most studies to date focus on molecular controls over diversity of neuron subtypes and broad patterns of connectivity– further focusing on transcriptional regulators in nuclei, and gene expression in somata/nuclei– increasing evidence indicates that local molecular controls in GCs (and thus developing and mature synapses) are critical in establishment and function of distinct cortical circuits.

Here, we focus investigation on functions of the highly penetrant, monogenic ASD- and ID-risk gene *Bcl11a* in CPN_SL_ circuitry as an exemplar of subcellular localization of molecular machinery and its potential dysregulation more broadly. CPN transfer and integrate cortical information bilaterally, and thus function centrally in high-level associative, integrative, cognitive, behavioral, and sensori-motor functions. These fundamental and high-level functions rely on precise, area-specific CPN subtype connectivity and diversity. We apply multiple independent modes of investigation to elucidate the requirement for proper *Bcl11a* function in area-specificity and precision of interhemispheric connectivity, associative social behavior, and subcellular localization of molecular machinery regulating precision of connectivity. Our experiments identify that: 1) disrupting proper subcellular localization of GC molecular machinery *in vivo* causes aberrant axonal targeting and circuit formation that persists into adulthood, affecting very specific social and cognitive behaviors; 2) recently developed *in vivo* experimental and analytical approaches enable identification of a focused set of dysregulated molecular controls in *Bcl11a* null GCs, critical for precise axonal targeting and otherwise not detected by investigating only somata; 3) in some cases, soma-only investigation would mistakenly detect the opposite direction of dysregulation from that at the functional GC, since some regulation of subcellular molecular machinery is likely due to differential trafficking, rather than being transcriptional. Together, the results presented here reveal multiple interacting layers of subcellular molecular controls functioning importantly in development of precise and highly disease-relevant cortical circuitry, function, and behavior, which would otherwise remain unidentified or misidentified by investigation of somata only.

### Bcl11a expression in CPN_SL_ is essential for precise targeting and refinement of connectivity between the two hemispheres

Using these multiple investigative approaches, we identify that deletion of *Bcl11a* from CPN_SL_ cell-autonomously causes striking disruption of appropriate and precise areal targeting. This aberrant circuit formation perdures into adulthood and is at least in part due to significant changes in subtype-specific subcellular GC molecular machinery, which we find to be critical for precise contralateral homotopic CPN_SL_ targeting. Importantly, deletion of *Bcl11a* from CPN_SL_ does not disrupt axonal elongation or midline crossing. We previously identified that wider forebrain-specific deletion of *Bcl11a* causes agenesis of the CC with prominent Probst bundles, which contain axons that failed to cross the midline^12^. This abnormality is likely due to fate shift of many cingulate cortex CPN into subcerebral PN, thus failing to properly “pioneer” the CC, since cingulate axons are the earliest population to cross the cortical midline. Since our experiments here delete *Bcl11a* only from CPN_SL_, our results indicate that *Bcl11a* null CPN_SL_ are able to effectively fasciculate and follow the axonal tract pioneered by earlier midline crossing cingulate cortex CPN.

Intriguingly, we find that *Bcl11a* deletion causes an increased number of CPN_SL_ axons to cross the midline at P0, suggesting an increase in axon elongation speed (perhaps due to less precision of GCs evaluating cues along the trajectory) and/or axon number per neuron. Although we have observed no evidence that *Bcl11a* deletion increases the number of axons per neuron *in vivo*, which can occur in cultured hippocampal neurons^48^, we cannot absolutely exclude this possibility. However, normalizing the number of axons post-midline to pre-midline axon fascicles (Supp. Fig. 1g) supports the interpretation that axon elongation speed is indeed increased. This interpretation is also supported by the observation that mitochondria– key in axon extension– are dysregulated in *Bcl11a* null CPN_SL_ GCs (see below) and is consistent with increased axon dynamics and outgrowth in cultured hippocampal neurons following *Bcl11a* knockdown^48^. Accelerated axon elongation, potentially caused by disrupted subcellular GC molecular machinery (Fig. 3), likely contributes to aberrant contralateral area-specific targeting by CPN^SL^ axons following CPN-specific *Bcl11a* deletion.

The CC and AC are the two dominant forebrain interhemispheric axonal tracts connecting between regions of the two hemispheres^35^. The AC is present in all vertebrates and normally is composed of connections from the lateral neocortex and piriform cortex^4^, as well as olfactory and amygdaloid regions^35^, whereas the CC more recently emerged in eutherians and contains the overwhelming majority of interhemispheric connections from evolutionary newer CPN^SL.^ The population of interhemispheric connections (in particular, by CPN) expanded dramatically in mammals, especially between sensorimotor cortical regions. Disruption of the AC is associated with neurodevelopmental and neuropsychiatric disorders, including ASD^49^, and rerouting of cortical connections via the AC is observed in human agenesis of the CC^50,51^. We previously identified that forebrain-specific deletion of *Bcl11a* causes enlargement of the AC, with an increase in somatosensory cortical neurons projecting through this tract^12^. Here, we build on those results to identify that loss of *Bcl11a* function from only CPN_SL_ cell-autonomously causes aberrant projections through the AC (Fig. 2). These results suggest that *Bcl11a*, in coordination with other CPN_SL_ molecular controls, enabled both expansion of evolutionarily newer CPN_SL_, as well as increased precision of contralateral targeting by re-routing CPN_SL_ axonal projections from the AC to the CC. The results presented here indicate that deletion of *Bcl11a* causes “reversion” of CPN_SL_ to an alternate, evolutionarily older, interhemispheric projection neuron identity. In support of this interpretation, deletion of *Satb2*, another important molecular control over CPN subtype development, also causes re-routing of callosal axon projections via the AC in mice^52^. Together, these results suggest that Bcl11a and Satb2 might interact in CPN_SL_ to regulate axonal projection and development of cortical neuron identity, in particular for CPN.

Perhaps the most strikingly aberrant projection by *Bcl11a* null sensorimotor CPN_SL_ is a seemingly highly specific projection to the BLA, essentially absent in WT brains (Fig. 2). The BLA normally receives multiple inputs from various brain regions including prefrontal, piriform, auditory, associative, and insular cortices^53,54^. However, unlike in primates, the BLA does not normally receive any direct input from primary sensory cortex in rodents. Disruption of BLA function is frequently associated with neuropsychiatric and neurodevelopmental disorders, including ASD^55^. Importantly, BLA has been shown to be important for normal social behavior, cognition, and anxiety^56,57^. Therefore, aberrant sensorimotor CPN_SL_ projections to the BLA (bilaterally, with *Bcl11a* deletion from only unilateral CPN_SL_), in addition to disruption of precise contralateral CPN targeting, might likely together contribute importantly to defects in social novelty preference and working memory in *Bcl11a* cKO mice (Fig. 3). How *Bcl11a* null sensorimotor CPN_SL_ axons target the BLA so specifically, and bilaterally, is both fascinating and entirely unknown. The ipsilateral projection appears quite direct and targeted. Since the AC relays reciprocal connections between the two amygdalae, it is possible that some CPN_SL_ axons project further, through the AC, to reach the contralateral amygdala^49^. Future investigation of subcellular GC and soma transcriptomes of *Bcl11a* null, BLA-projecting CPN_SL_ have potential to identify local molecular controls underlying these remarkably aberrant and clinically relevant projections.

### Expression of Bcl11a by CPN_SL_ during development is necessary for normal adult behavior

People with *BCL11A* mutations/deletions exhibit behavioral deficits, including abnormal social interaction, intellectual disability, repetitive behavior, and anxiety^15^. Our results demonstrate that even spatially restricted loss of *Bcl11a* from a relatively modest subset of CPN_SL_ also produces homologous ASD-associated behavioral deficits in mice, likely due to the observed abnormalities of CPN_SL_ projections. Consistent with this idea, we observed specific abnormalities of social novelty preference and working memory in *Bcl11a* null CPN_SL_ mice, rather than widespread disruption of behavior. Although we cannot unequivocally determine whether deficits in social novelty preference stem from a primary lack of interest in social novelty, and/or from the deficit in working memory exhibited by *Bcl11a* cKO mice, this end behavior reveals that correct CPN_SL_ circuit development is critical for social novelty preference, without an equivalent level function in general mouse-to-mouse sociability or other commonly investigated cognitive and/or ASD-associated behaviors.

Though we employed more specific deletion of *Bcl11a* from only a spatially delineated subset of CPN_SL_ in otherwise WT mice, our results are consistent with those of Dias *et al*. (2016)^15^, who reported that *Bcl11a* (full-body) heterozygous mice display abnormal social novelty preference/discrimination. The same study also reported broader disruption of sociability behavior by these mice. This difference from our more specific findings of disruption of social novelty preference is likely due to the fact that *Bcl11a* is expressed by many other neuronal populations with much broader behaviorally relevant circuitry, including CThPN and subsets of cortical interneurons^32^, midbrain dopaminergic neurons^58^, and hippocampal neurons. These non-CPN populations function in a range of broader sensori-motor gating, multi-system inhibition, reward and motor system tone, learning and memory, and are all spared by the subtype- and spatially-specific manipulations we employed. Further, though neuronal birthdate-timed *in utero* electroporation enabled efficient, widespread targeting of sensorimotor area CPN_SL_, a subset of CPN in the targeted area will evade labeling and remain wildtype. Importantly, both our results and Dias *et al*. identify that *Bcl11a* expression during brain development is necessary for normal social novelty preference/discrimination.

### Loss of Bcl11a function causes substantial and selective dysregulation of subtype-specific CPN_SL_ subcellular GC transcriptomes during development

How function-specific cortical circuitry is regulated with specificity and precision beyond nuclear transcriptional controls is still an underexplored question, partially due to previous lack of applicable subtype- and stage-specific subcellular approaches. Here, we identify that subtype-specific deletion of *Bcl11a*, a transcriptional regulator, causes disruption of circuit-specific axon GC transcriptomes (Fig. 4), with changes in developmentally and subcellularly functional transcripts likely leading to disruption of CPN_SL_ axonal connectivity, targeting, function, and ultimately behavior of adult mice. Loss of *Bcl11a* appears to dysregulate the CPN_SL_ subcellular GC transcriptome in multiple ways, both expected and surprising: 1) as expected, *Bcl11a* loss of function secondarily dysregulates the abundance of some GC transcripts due to differential somatic expression, seemingly driven by soma transcription levels; 2) increasing the abundance of many normally GC-enriched transcripts, decrease the abundance of many normally soma-enriched transcripts in CPN GCs; and 3) surprisingly and quite intriguingly, Bcl11a appears to specifically regulate trafficking of a subset of transcripts toward, or excluding from, GCs, despite negligible changes in soma-localized abundance of the same transcript(s), or even with soma-localized changes in the opposite direction (Fig. 3b-c).

There are multiple potential molecular mechanisms that might dysregulate trafficking and/or localization of specific transcripts to axon GCs in *Bcl11a* null CPN. One potential mechanism is dysregulation of RNA binding proteins (RBPs) that are critical for RNA trafficking to GCs^59^. Interestingly, we identified eight RBPs that are differentially expressed in *Bcl11a* null CPN_SL_ somata (Cirbp, Enox1, Khdrbs2, Qk, Rbm24/38, Sf3b4, Tia1), while no RBP RNAs are differentially abundant in GCs, consistent with soma-to-GC RBP functions as proteins. Intriguingly, 3’ UTR motifs that are bound by the RBPs Qk and Rbm24/38 are underrepresented in enriched DAGs, and a 3’ UTR motif that is bound by the RBP Nova2 is overrepresented in depleted DAGs^60^. While we did not detect *Nova2* transcripts, two of its interactors (*Srrm4* and *Fam107b*) are significantly increased in *Bcl11a* null somata, as are *Qk* and *Rbm24/38*^61^.

These data raise the possibility that binding of these RBPs to transcripts prevents or reduces trafficking to GCs, and that deletion of *Bcl11a* leads to increased somatic expression of these RBPs, preventing certain transcripts from being trafficked to GCs at appropriate levels.

Quite interestingly, we also observe broad enrichment of mitochondria-related GO terms among *Bcl11a* DAGs (Fig. 4, Supp. Fig. 9e). Mitochondrial dysfunction is frequently associated with neurodevelopmental disorders. 7q11.23 deletion in William syndrome, a rare genetic developmental disorder characterized by mild to moderate ID, dysmorphic facial features, and disrupted metabolism, reduces copy number of DNAJC30, a protein chaperone that functions in stability of OXPHOS supercomplexes. Deleting *Dnajc30* in mouse disrupts mitochondrial function in neocortical neurons and Dnajc30-/-mice exhibit thinner callosal axons, aberrant dendritic arborization, and disrupted social and anxiety-like behaviors^62^. It is also well-established that mitochondria play important roles in multiple stages of axonal development, including axon extension and branching^63–65^. Because axon extension is energetically demanding, dynamic and appropriate regulation of GC-localized mitochondria is key^66^. Axon GCs contain higher densities of mitochondria during extension, and blocking extension results in redistribution of mitochondria away from GCs^67^. However, it is likely that mitochondrial function is required primarily for general aspects of axon development, and is less critical for precise, circuit-specific axonal targeting. Although it is currently not clear whether and how changes in mitochondria-related genes in *Bcl11a* null CPN_SL_ affect mitochondrial number or function, it is possible that dysregulation of mitochondrial function in GCs contributes to aberrant axon elongation of null CPN_SL_.

Importantly, *Bcl11a* deletion does not nonspecifically dysregulate core GC subcellular transcripts, important for generic axonal motility, dynamics, and/or extension, but instead causes specific dysregulation of transcripts whose protein products are likely important for precise CPN targeting (Figs. 4 and 5). We functionally investigate two of these transcripts: *Mmp24* and *PcdhαC2*, and identify that each is necessary for precise contralateral targeting of CPN_SL_ axons (Fig. 4). *Mmp24* encodes a member of the membrane-type matrix metalloproteinase family that cleaves most ECM components on cell surfaces^43^. Consistent with data from other groups showing that Mmp24 localizes to GCs of cultured DRG and hippocampal neurons, and regulates axonal growth^43^, we find that Mmp24 localizes to axon GCs of cultured CPN_SL_ (data not shown), and that specific *Mmp24* reduction in CPN_SL_ causes aberrant medial and lateral targeting (Fig. 5). Together, these results suggest that Mmp24 likely participates in CPN axon outgrowth, guidance, and/or homotopic targeting, potentially by regulating cleavage of receptors, ligands, and/or ECM components.

*PcdhαC2* is another transcript dysregulated in *Bcl11a* null CPN_SL_ GCs. *PcdhαC2* is a member of the protocadherin family, which function as molecular “barcodes” for self-recognition/avoidance, tiling^68^, and establishment of subtype-specific circuits^68–70^. PcdhαC2 is a member C-type Pcdhs, which are characterized by constitutive expression in Purkinje neurons^69,70^ and exert specific “carrier” function by transporting Pcdhα isoforms to cell surfaces^71^. PcdhαC2 is also required for axonal tiling and assembly of serotonergic circuitry^69^, and has been identified as an ASD-risk gene in the SFARI human gene dataset. High specificity is key in homophilic interaction of Pcdhs, since absence of the correct, specific Pcdh prevents proper binding^71^. Consistent with these findings, our data reveal that PcdhαC2 functions in precision and spatial organization of homotopic contralateral targeting by CPN_SL_ *in vivo*, perhaps contributing to formation of unique “barcode(s)” critical for precise axonal targeting.

Our approaches appear to enable identification of molecular machinery that functions in alternate axons of the same subtype (e.g. somatosensory area CPN_SL_), or potentially of the same neurons (some subpopulations of CPN have at least two axons)^72,73^. Unlike *Bcl11a* deletion from CPN_SL_, knockdown of either *Mmp24* or *PcdhαC2* from CPN^SL^ does not cause aberrant projections of CPN^SL^ to the amygdala. Since we purified GCs and somata from, and investigated subcellular transcriptomes of, contralaterally targeting CPN_SL_, it is highly likely that these GCs do not contain transcripts that would cause amygdala targeting. Future studies could investigate subcellular transcriptomes of amygdala-projecting GCs from *Bcl11a* null CPN_SL_, potentially enabling identification of distinct subcellular targeting mechanisms that are aberrantly engaged with subtype-specific deletion of *Bcl11a* from CPN_SL_. Both sets of aberrant subcellular GC machinery are likely highly relevant to elucidating mechanisms of developmental cortical circuit dysgenesis and behavioral changes in ASD.

## Methods

### DNA constructs

Cre-dependent tdTomato and EGFP constructs were derived by subcloning into a pCBIG vector backbone, a gift from C. Lois. A myristoylation sequence was inserted to the N-terminus of the fluorophore sequences to achieve membrane targeting. The coding sequence for CreM was obtained from Addgene (Kaczmarczyk and Green JE, 2001; Addgene plasmid #8395). These constructs were used in Figures 1 and 2 and associated supporting figures. Constitutively expressing myristoylated-EGFP and tdTomato constructs were cloned into the pCBIG vector backbone, and used in Figures 3, 5, and associated supporting figures. For GC and soma purification by FSPS and FACS, respectively, in Figure 4 and supporting figures, a 2A bi-cistronic construct encoding myristoylated-tdTomato and Histone2B-GFP, or a 2A bi-cistronic construct encoding myristoylated-EGFP and Histone2B-tdTomato were cloned into the pCAG vector backbone. Control non-targeting shRNA (Sigma, SHC002), and shRNAs targeting *Mmp24* and *Pcdhαc2* were obtained from Sigma (*Mmp24* shRNA, Clone ID: NM_010808, TRC Number: TRCN0000033008, Target sequence: GCCCGCATAGACGCAGCCTAT; *PcdhaC2* shRNA, Clone ID: NM_001003672.1-2310s1c1, TRC Number:

TRCN0000094942, Target sequence: CCTCAAAGTACAGCCTCACTT). For knockdown with CRISPR-CasRx, pXR001 2A bi-cistronic construct encoding EF1a promoter driven *Cas13d* gene and EGFP was used to express *Cas13d*. pXR001: EF1a-CasRX-2A-EGFP was a gift from Patrick Hsu (Addgene plasmid # 109049; RRID:Addgene_109049^74^). CasRX compatible guide RNA expression was achieved by cloning the guide sequences into the pXR003 vector backbone (pXR003: CasRx gRNA cloning backbone was a gift from Patrick Hsu (Addgene plasmid # 109053; RRID:Addgene_109053). Target specific CasRX guide RNA sequences are as follows: *Mmp24* guide RNA #1: AGCTTTCAAGAAGTGAGCACCAA; *Mmp24* guide RNA #2: GCCACACATCGAAAGCCTGACGA; *Mmp24* guide RNA #3: AAGATAGTGTAGAGCAGCACCAG; *PcdhαC2* guide RNA #1: AGAACCAAGTAGTGCAAGGCAGT; *PcdhαC2* guide RNA #2: TAAATGGTAAAAGAGCCAAGCAT; *PcdhαC2* guide RNA #3: TGTATAAGTCTGTGAGCACCACC.

### Mice

All mouse work was approved by the Harvard University Institutional Animal Care and Use Committee, and was performed in accordance with institutional and federal guidelines. Outbred strain CD1 mouse pups used for some *in utero* electroporation experiments were ordered from Charles River Laboratories (Wilmington, MA). *Bcl11a*^*fl/fl*^ mice were generated and generously shared by H. Tucker, G. Ippolito, and colleagues (RRID: MGI _4358088, Sankaran et al., 2009; Lee et al., 2013). All *Bcl11a*^*fl/fl*^ and *Bcl11a*^*fl/+*^ mice were maintained on a CD1 or C57BL/6J background. The morning of vaginal plug identification was designated as E0.5, and the day of birth as P0. Mice were housed in groups of 2-4, except males that were single housed post-breeding for timed pregnancy, all on standard 12h light / 12h dark cycle.

### Immunocytochemistry and histology

Mice were transcardially perfused with 4% paraformaldehyde (PFA). Following dissection, brains were postfixed overnight in 4% PFA, then transferred to 30% sucrose/PBS solution for cryoprotection at 4 °C. Tissues were embedded in O.C.T. compound (Electron Microscopy Sciences), and sectioned at 50 µm sections using a cryostat (Leica). Sections were stored in PBS with 0.05% sodium azide solution for later experiments. Sections were blocked for 1-2 h with blocking solution (1x PBS + 10% goat normal serum + 0.5% Triton-X). Following blocking, sections were incubated with primary antibodies overnight at 4 °C in blocking solution. Incubation with secondary antibodies was performed for 2 h at room temperature. Finally, sections were mounted using Fluoromount-g with DAPI (Southern Biotech). Images were acquired on epifluorescence microscope (Nikon 90i).

### Western blot analysis

GCF samples were lysed and protein concentration was determined using the fluorometric Qubit protein assay. Equal amount of total protein was loaded for each sample. Protein samples were run on 4-20% precast polyacrylamide gel (4-20% Mini-PROTEAN TGX Precast Protein Gels, BioRad) to separate them and transferred onto PVDF membranes using semi-dry transfer. Membranes were blocked using 3% bovine serum albumin prepared in TBS-T (50 mM Tris-HCl (pH 7.4), 150 mM NaCl, 0.2% Tween-20) for 1 hour at room temperature. Membranes were incubated with primary antibodies overnight at 4 °C. Next, they were washed three times with TBS-T, followed by incubation with horseradish peroxide-conjugated secondary antibodies (Life Technologies; Abcam) for 1 hour. Following washing with TBS-T, immunoreactivity signals were visualized through detection of chemiluminescence by SuperSignal West Pico PLUS (ThermoFisher) using a CCD camera imager (FluoroChemM, Protein Simple).

### Antibodies

The following primary antibodies and dilutions were used: rabbit anti-RFP, 1:2000 (Rockland Immunochemicals, Catalog no. 600-401-579); chicken anti-GFP, 1:1000 (Thermo Fisher, Catalog no. A10262), rabbit anti-Mmp24, 1:250 (Abcam, Catalog no. ab233044). Alexa Fluor-conjugated secondary antibodies (Invitrogen) were used at a dilution of 1:500.

### *In utero* electroporation

The Institutional Animal Care and Use Committee of Harvard University approved all experiments and procedures. Following anesthetization of timed pregnant mice, the uterine horns were exposed, and the plasmids mixed with Fast Green dye (Sigma) were microinjected into one of the lateral ventricles for CPN circuit analysis and axon GC isolation experiments. Five 34 volt pulses (50 ms on and 950 ms off per pulse) were delivered in appropriate orientation across the embryonic head using 5mm diameter platinum plate electrodes (CUY650-P5, Protech International Inc) and a CUY21EDIT square wave electroporator (Nepa Gene, Japan). For behavioral experiments, bilateral targeting was achieved by simultaneously microinjecting the plasmids into both lateral ventricles, followed by delivery of current while electrodes were placed in the middle of both hemispheres. In the DNA mixture, the fluorophore-expressing plasmid concentration was around 2 µg/µl for CPN circuit analysis, 3 µg/µl for behavioral experiments, 4 µg/µl for GC and soma sort experiments, 0.5 µg/µl for the Cre-expression plasmid, and 3 µg/µl for shRNA plasmids. For dual population mosaic labeling^34^, the plasmid concentrations were as follows: 30-50 ng/µl for the Cre-expression plasmid, 0.5 µg/µl for the CreM plasmid (Cre-inducible Cre recombinase), 2 µg/µl for the Cre-activated fluorophore, and 2 µg/µl for the Cre-silenced fluorophore. For CRISPR-CasRX experiments, the ratio between *Cas13d* gene expression and mix of three guide RNA constructs was 2:3 in co-expression DNA mixture.

### Quantification of sensorimotor superficial callosal connectivity

*For dual-population mosaic analysis (Figures 1 and 2)*: Entire coronal brain sections containing interspersed control and *Bcl11a* null/heterozygous null populations were imaged for GFP and tdTomato. *For non-mosaic analysis (Figure 5):* Entire coronal brain sections from each experimental condition, all with matching electroporated ipsilateral targeting, were imaged for tdTomato. To quantify fluorescence intensity across the entire mediolateral extent of cortex (from midline to the end of insular/piriform cortex), images were warped into a rectangle shape using perspective warping function of Photoshop. The resulting rectangular image was thresholded, then binned into 400 segments using internally developed ImageJ macro called “*Make400ROI* and *measure*.*jim”*. The fluorescence intensity in each bin was measured using the same macro. The fluorescence intensity in each bin was normalized to the total fluorescence intensity of 400 bins. A plot of normalized average fluorescence intensity (and S.E.M.) across the mediolateral positions was generated.

### qRT-PCR

Total RNA was collected using RNeasy Plus kits (Qiagen). Reverse transcription of mRNA transcripts to produce cDNA for qPCR was achieved using Superscript IV Reverse Transcriptase (Invitrogen). qPCR was performed using SsoAdvanced Universal SYBR Green Supermix (Bio-Rad) on a QuantStudio 3 Real-Time PCR System (ThermoFisher Scientific). Reactions were run in triplicate, and averages of these triplicates were used for statistical analysis. *Gapdh* was used as an internal control. Primer sequences used for qRT-PCR are as follows: *Gapdh* (NM_008084), Forward sequence: CTTTGTCAAGCTCATTTCCTGG, Reverse sequence: TCTTGCTCAGTGTCCTTGC; *Mmp24* (NM_010808), Forward sequence: GATTCAGATGAACCCTGGACG, Reverse sequence: CATGTATTGGTAGAAGGGAGCC; *PcdhαC2* (NM_001003672), Forward sequence: GTACCGGGAACCTGATTATCC, Reverse sequence: GACCAGCCCGTAGAATGC.

### GC fractionation and purification (FSPS)

Fluorescently labeled CPN_SL_ axonal GCs were fractionated and isolated as previously described in detail elsewhere^32,33^. Briefly, 5-10 cortical hemispheres containing axonal GCs were micro-dissected rapidly and homogenized in 0.32M sucrose supplemented with 4mM HEPES (Thermo Fisher, 15630106), Halt protease and phosphatase inhibitor cocktail (Thermo Fisher, 78442), and RNasin ribonuclease inhibitors (Promega, N2515) at 900 RPM in a glass-Teflon tissue grinder with 10-12 strokes. Following a low speed centrifuge at 1700 RPM for 15 min at 4 °C, “post-nuclear homogenate” (PNH; input) supernatant was collected, and layered onto 0.83 M and 2.5 M sucrose gradient (Top-to-bottom: Input, 0.83 M sucrose, 2.5 M sucrose). The discontinuous sucrose gradient was centrifuged at 242K g for 47 min at 4 °C in a vertical rotor (VTi50 rotor, Beckman Coulter). The bulk GC fraction (GCF) was extracted from the interface between 0.32 M and 0.83 M sucrose.

Labeled CPN_SL_ axonal GCs were purified from bulk GCF via fluorescent small particle sorting (FSPS) utilizing customized BD SORP FACS AriaII instrument^32,33^. Bulk GCF in sucrose solution were diluted with 4-to 6-fold in chilled PBS prior to loading into the sorter. GCs were sorted directly into the RLT lysis buffer (QIAGEN Allprep Mini Kit) for total RNA extraction.

### Soma Isolation (FACS)

Fluorescently labeled CPN_SL_ somata were isolated according to established approaches^33^. Briefly, *in utero* electroporated cortical hemispheres were collected in chilled dissociation solution and micro-dissected in chilled HBSS using a fluorescent dissection microscope to remove meninges and ventricular zone, and enrich for labeled CPN. Cortical tissues were returned to dissociation solution and enzymatically dissociated for FACS. Fluorescently labeled somata were isolated from bulk soma prep by sorting on a FACS AriaII instrument equipped with and 85 µm nozzle operated at 45 p.s.i. Somata were sorted directly into the RLT lysis buffer (QIAGEN Allprep Mini Kit) for total RNA extraction.

### Genome-wide RNA-sequencing and bioinformatic analyses

Following total RNA extraction from purified GCs and somata, RNA was quality controlled and quantified using an Agilent 2100 Bioanalyzer. Next, purified RNA samples were converted to cDNA using the SMART-seq v4 Ultra Low Input RNA Kit (Takara Bio), and cDNA libraries were generated with the Nextera XT DNA Library Preparation Kit (Takara Bio) by the Harvard University Bauer Core Facility. High-throughput sequencing was performed by loading the pooled GC and soma cDNA libraries on an Illumina Hiseq 2500 Rapid Run platform. The raw FASTQ files of 75 bp paired-end reads were collected for downstream analysis.

Adapter sequences were clipped, and unpaired reads were trimmed with Trimmomatic version 0.39. Reads were aligned to the GRCm39 mouse genome with STAR in 2-pass mode^75^, and reads mapping to exons were quantified with featureCounts^76^. To identify “ambient” genes that arose from supplementing labeled B6 cortices with unlabeled CD1 hemispheres during GC purification, variants were called on all samples with GATK HapotypeCaller^77^. 34,743 genes were retained for differential analysis, since their CD1 penetrance was significantly lower (q < 0.0001, one-sided binomial test) than the mass percentage of CD1 tissue input in at least four of the six GC samples, while 2,364 genes did not pass this quality control test and were not evaluated for differential expression. Differentially expressed genes between WT GCs and *Bcl11a* null GCs were identified with DESeq2, using an FDR of 0.05^78^. 13,487 genes had a normalized mean count of >= 32, the independent filtering threshold, and effect sizes were shrunken via the apeglm method^79^. Prior to clustering, counts were transformed with the regularized algorithm^78^. Similar steps were followed for all differential expression analyses. GO enrichment analysis was performed with ClusterProfiler, and terms were reduced based on semantic similarity with Revigo. The data discussed in this publication have been deposited in NCBI’s Gene Expression Omnibus and are accessible through GEO Series accession number GSE211405 (https://www.ncbi.nlm.nih.gov/geo/query/acc.cgi?acc=GSE211405). All code is available on GitHub (https://github.com/detillman/bcl11a_macklis_2022).

### Behavioral Experiments

For behavioral analyses of *Bcl11a* CPN cKO and control mice, experiments were started at 8 weeks of age. All mice were gently handled for 3 days before each experiment to acclimate the mice to the experimenter, and to reduce stress on the day of testing. All behavioral experiments were performed during the light cycle. The behavioral experimenter was blinded to the group allocation of each mouse both during the experiment and when assessing the behaviors and results. Ear-tag identification numbers were used until outcome analyses were finalized. Mice that were tested in multiple behavioral paradigms were given a minimum of 5 days resting period between experiments. Mice without successful bilateral CPN targeting (e.g. GFP+ cells in only one hemisphere) were excluded from the study.

#### Open Field Test

Mice were individually placed in an open field arena (40 cm square arena with 30 cm height) and the activity of each mouse was monitored and video recorded over a 10-minute period. Videos were processed and analyzed with Ethovision XT 14 software (Noldus, VA) to assess total distance moved, velocity, total time in motion, time spent in the center of the arena, and other basic locomotion parameters.

#### Light-Dark Exploration Test

The light-dark apparatus (40 cm x 40 cm) consisting of two chambers – a light chamber (20 cm x 20 cm) covering half of the arena, and a black box (20 cm x 40 cm) covering the other half with a 7.5 cm x 7.5 cm door between the two chambers through which a mouse can pass – was used for light/dark exploration testing. For testing, initially the door between the light and the dark chambers was closed, and a test mouse placed in the dark chamber. Following a brief acclimation period, the door between the chambers was removed, and the mouse was allowed to explore the arena for 10 minutes, moving freely between both chambers. The activity of each mouse was monitored and video recorded, and the videos were analyzed later with Ethovision XT 14 software (Noldus, VA) to assess the time spent in each chamber, latency to enter light chamber, and the number of visit to the light chamber.

#### Elevated Plus Maze

The elevated plus maze employed here consists of a black acrylic maze with four white floor arms, two of which are open without walls, and two are enclosed. Each arm is 26 cm x 6 cm, with 3 cm high closed arm walls. The maze is elevated 40 cm from the floor. For testing, individual mice were placed at the junction of the open and closed arms, and allowed to explore the arena freely for 10 minutes. The activity of mice was monitored and video recorded, and the videos were analyzed later with Ethovision XT 14 software (Noldus, VA) to measure the time spent on each arm, and the number of entries to each arm. An entry was defined as a mouse having the front paws and the half of body on any of maze’s arms. The 6 cm x 6 cm junction area was not considered a part of any of the arms.

#### Three-chamber Sociability and Social Novelty

The three-chamber social interaction test chamber employed here consists of a rectangular black acrylic box (60 cm x 40 cm) with 40 cm height. The arena is split into three chambers, each 20 cm wide, separated by black acrylic. Each inner wall has a 7.5 cm x 7.5 door that can be sealed. The wire cages placed in the side chambers are cylindrical with a bottom diameter of 10 cm, with the bars were spaced 1 cm apart to allow test animals to see and interact with stranger mice. Age and sex matched C57/Bl6 mice, housed prior to testing in cages separate from the test mice on separate racks, were used as novel stranger mice. Stranger mice were acclimated to the wire cages and the maze a day before testing to reduce stress during the assay. On the day of testing, test mice were placed in the middle chamber of the arena, and given 5 seconds to acclimate with doors to side chambers closed. Then, the doors were opened, and the test mouse was given 10 minutes to habituate/explore the empty arena with empty wire cages in place. Following the 10 minutes habituation period, each test mouse was guided back to the middle chamber, and the doors were closed. An age and sex matched novel mouse (Stranger 1) was placed in one of the wire cages, and an inanimate object was placed in the other. Each test mouse was again given 10 minutes to explore all three chambers freely. Following the second trial, each test mouse was again brought back to the middle chamber, and the inanimate object was replaced with a second novel mouse (Stranger 2). Each test mouse was allowed to explore all three chambers freely for an additional 10 minutes. The activity of mice during each trial was monitored and video recorded, and the videos were analyzed later with Ethovision XT 14 software (Noldus, VA) to measure time spent in each chamber, and time spent in close interaction with the wired cages (within 5 cms and facing the wired cages).

#### Novel Object Location and Recognition

The apparatus employed here consists of a 40 cm square black acrylic box with 30 cm height (open field box). The novel object location and recognition chamber is fitted with visual cues on each wall to provide adequate spatial discrimination. Additional visual cues placed in the behavioral test room walls. Two identical objects and a distinct object with similar size are used as “novel objects” for mice to interact with. Behavioral testing consists of 2 days of 5-minute habituation to the arena without any objects, followed by an additional 5-minute habituation on the day of novel location testing. Then, two identical objects were placed close to one of the walls, and each test mouse was allowed to explore the arena for 5 min (“training”). For novel location testing, one of the identical objects was moved close to the opposite wall, and each mouse was allowed to explore the arena and the objects for 5 minutes. Next day, one of the objects is replaced with the different “novel object”, and each mouse was again allowed to explore the arena and objects for 5 minutes. The activity of each test mouse was monitored and video recorded, and the videos were analyzed later to assess time spent interacting with each object.

#### Spontaneous Alternation Test (Y-Maze)

The symmetrical Y-maze employed here apparatus has a white acrylic bottom plate with black walls. Each arm of the Y-maze is 30 cm long, 10 cm wide, and the walls are 20 cm high. The walls at the end of each arm were marked with a distinct black and white pattern (e.g. vertical or horizontal stripes) to provide guidance cues. The test consisted of a single 10-minute trial, during which each test mouse was allowed to explore all three arms of the Y-maze freely. Each mouse was placed at the end of one arm. The arms were labeled as “A”, “B”, and “C”. The “start” arm was varied between mice to avoid placement bias, alternating randomly between A, B, and C. The activity of mice was monitored and video recorded, and the videos were analyzed later with Ethovision XT 14 software (Noldus, VA) to assess the sequence of arm entry. Spontaneous Alternation [%] was defined as consecutive entries to 3 distinct arms (e.g. A, then B, then C, or A, then C, then B), divided by the number of possible alternations (total arm entries-2).

#### Barnes Maze

The Barnes maze is widely used to test rodent spatial memory, both long- and short-term. The Barnes maze employed here consists of an 80 cm diameter circular white acrylic, 1.3 cm thick. The maze has 20 holes of 5.5 cm diameter around the perimeter, each 2.5 cm from the edge of the maze. The platform is mounted on 40 cm legs. Additionally, the maze contains an acrylic escape box fitted below one of the holes (“target”), and a glass beaker that serves as the “start chamber”, allowing test mice to observe the maze and visual cues. Each wall of the test room was fitted with a distinct marking (e.g. a black poster with white vertical lines) to provide visual cues. On day 1 of the assay, each mouse was trained on the maze. Habituation included 30 seconds in the start chamber, followed by 15 seconds of manual guiding of each test mouse in the start chamber to the “target”. Each mouse was given three minutes to enter the escape box on its own, after which it was guided into the escape box. Each test mouse was given one minute in the escape box with the glass beaker occluding the hole (bottom down) to prevent exit during this period. On days 2-5, each test mouse was placed in the start chamber for 30 seconds, after which the start chamber was removed, and the mouse was given three minutes to locate the target and enter the escape box. In the event that a mouse did not find or enter the escape box, the experimenter gently guided the mouse into the escape box using the start chamber. Each test mouse was given one minute in the escape box. This training was repeated two more times on the same day, with at least 30 minutes between each training. On day 6 (probe test), the target/escape hole was sealed. Each test mouse was placed in the start chamber for 30 seconds, after which the start chamber was removed, and the mouse was given three minutes to explore the maze. On Day 13 (long-term probe testing), the target/escape hole was unsealed again. Each test mouse was placed in the start chamber for 30 seconds, after which the start chamber was removed, and the mouse was given three minutes to locate the target and enter the escape box. Each test mouse was given one minute in the escape box. The activity of mice was monitored and video recorded, and the videos were analyzed later with Ethovision XT 14 software (Noldus, VA) to assess latency to escape (days 3-5, and day 13), and time spent in each quadrant (i.e. target quadrant where the target/escape hole is located) on day 6 (probe test).

#### Marble Burying

The marble burying test is widely used to assess “compulsive-like” behaviors in rodents. The assay relies on the natural inclination to bury objects under stress and anxiety. The marble burying test employed here used standard rat cages (Allentown polycarbonate cage, 9.25 × 17.75 × 8 inches high) with filter-top covers. Cages were filled with 5 cm of standard corncob bedding used in home cages. 20 glass marbles, approximately 15 mm in diameter, were placed gently on the bedding surface in five rows of four, equidistant from each other and from the sides of the cage. Each test mouse was placed in the center of the prepared cage, and the cover was replaced. The mouse was given 30 minutes undisturbed to explore the cage. After 30 minutes, the test mouse was returned to the home cage, and the number of marbles buried was recorded. Scoring was done by two independent observers; averages of the independent scores were used for analysis. A marble was scored as buried if two-thirds of the surface area was covered by bedding.

### Statistics

All plots were generated and statistical analyses were performed using Graphpad Prism 9.0 software. Results are presented as mean ± s.e.m. Sample size was not predetermined but numbers of samples are consistent with established studies. Two-tailed *t*-tests were used for comparison of two data sets. One-way ANOVA followed by Bonferroni multiple comparison test was used for experiments with three or more data sets. Equal variances between groups and normal distribution of data were presumed, but not formally tested. Molecular and biochemical analyses were performed using a minimum of three biological replicates per condition. Behavioral experiments require larger data sets due to increased variability. A minimum of 6 animals per group was used for behavioral testing.

## Acknowledgements

O.D. and J.D.M conceived the overall project and experiments; O.D. and J.Y.K designed individual experiments, and OD., J.Y.K., D.T., Y.I., M.W., L.C.G, and T.A.A. performed the experiments; O.D., J.Y.K, D.T., and J.D.M. analyzed and interpreted the data, integrated the findings, and wrote and edited the manuscript. All authors contributed to discussions and manuscript editing.

**Acknowledgements**

We thank Megi Hoxha, Jaewon Heo, Prakruti Nanda, and Melody Ross for their technical support; members of the Macklis laboratory for scientific discussions and helpful suggestions; Joyce LaVecchio and Nema Kheradmand of the HSCRB-HSCI Flow Cytometry Core; the Harvard Center for Biological Imaging for infrastructure and support; Emma White, Christian Daly, and Claire Hartmenn of the Harvard Bauer Sequencing Core. Analytic tools and infrastructure in the Harvard Chan Bioinformatics Core were partially supported by funds from the Harvard NeuroDiscovery Center and the Harvard Stem Cell Institute. This work was supported by the following grants to J.D.M.: NIH DP1 NS106665; NIH R01 NS104055; NIH R21 NS104733; Simons Foundation Autism Research Initiative #515376; Paul G. Allen Frontiers Group— Allen Distinguished Investigator award #11855; Brain Research Foundation Scientific Innovations Award; Max and Anne Wien Professor of Life Sciences fund; Emily and Robert Pearlstein Fund; Jayne and Lee Seidman Fund; and additional infrastructure support from NIH grants NS045523, NS075672 and NS093376. O.D was partially supported by NIH Institutional Training Grant T32 AG000222. D.T. was partially supported by a 2022 NSF-Simon Center at Harvard University QBio Graduate Student Award. We dedicate this work to the memory of our friend and colleague John J. Hatch.

## Competing Interests

The authors declare no competing interests.

**Supplementary Figure 1:**
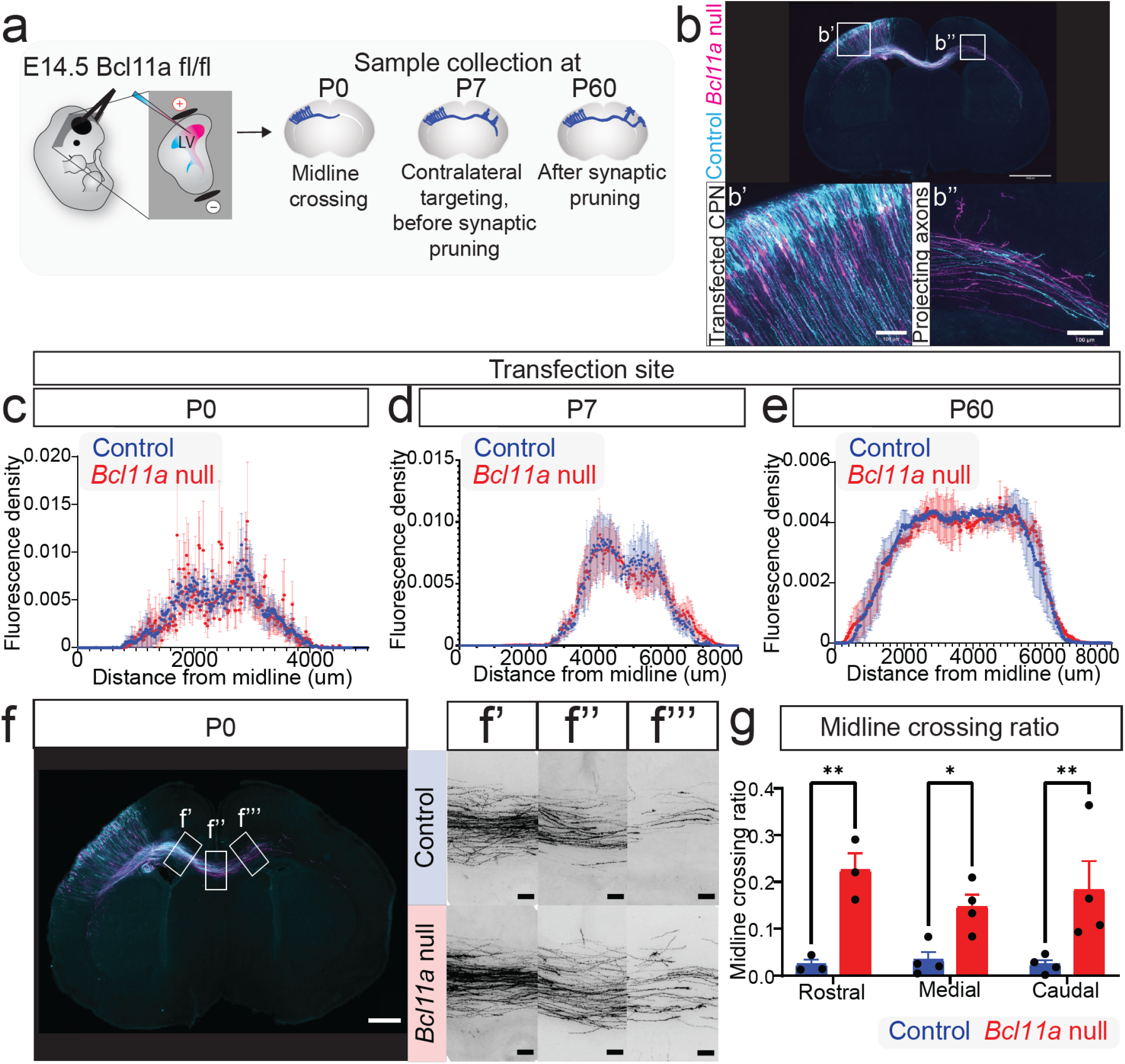
**a)** Schematic for analysis of CPN_SL_ connectivity. **(b)** Fluorescence images of P0 *Bcl11a*^flox/flox^ mouse brain electroporated *in utero* at E14.5 with a precisely-defined cocktail of Cre-recombinase, Cre-silenced myr-GFP expression construct (control; cyan), Cre-activated myr-tdTomato (*Bcl11a* null; magenta), and Cre-inducible Cre-recombinase. Two non-overlapping, genetically mosaic populations of interspersed control and *Bcl11a* null CPN_SL_ are generated. **(c-e)** Spatial distribution of *in utero* electroporation transfection regions of mosaic CPN_SL_ populations at P0 (c), P7 (d), and P60 (e) show highly equivalent transfection coverage between control and *Bcl11a* null CPN_SL_ somata (P0, n=6; P7, n=5; P60, n=3). **(f)** Fluorescence images of P0 *Bcl11a*^flox/flox^ mouse brain with mosaic populations of *Bcl11a* null (magenta) and control (cyan) reveals an increased number of *Bcl11a* null axons crossing the midline (f’’) and reaching the contralateral hemisphere (f’’’) compared to control, despite an equivalent number of pre-midline axonal fibers (f’). **(g)** Quantification of midline crossing ratio in rostral (motor), medial (somatosensory), and caudal (visual) cortex reveals that a higher percentage of *Bcl11a* null axons cross the midline compared to control (two-way ANOVA followed by Bo Bonferroni multiple comparison test; Control, n=3; *Bcl11a* null, n=3). All sections were immunolabeled for GFP (control; cyan) and RFP (*Bcl11a* null; magenta). Scale bars: 100 µm. *, p-value<0.033; **, p-value<0.0021. n, number of distinct brains used. Results are represented as mean ± s.e.m.

**Supplementary Figure 2:**
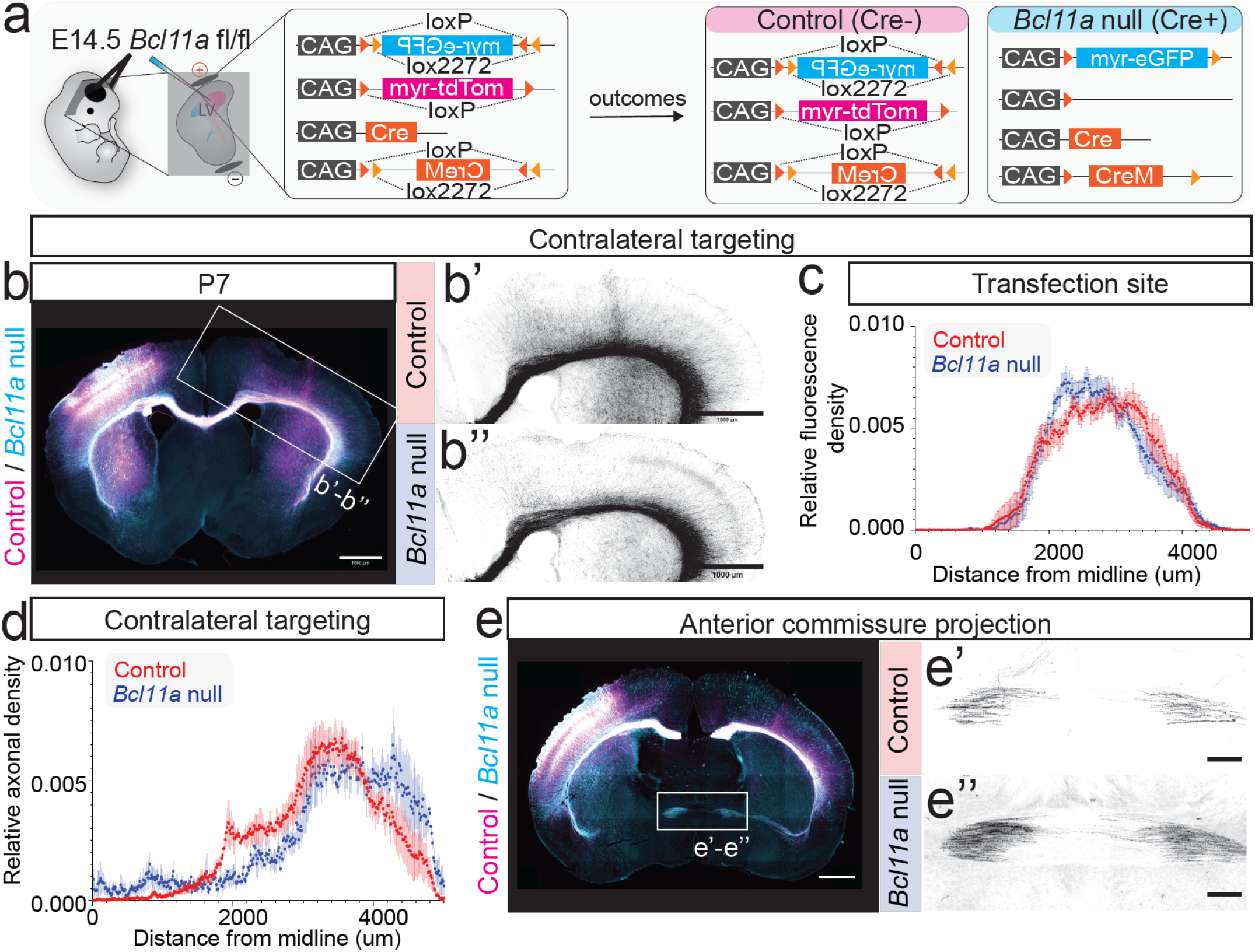
**(a)** Schematic for dual-population mosaic targeting of CPN_SL_ *in vivo. Bcl11a*^flox/flox^ mouse brain electroporated *in utero* at E14.5 with a precisely-defined cocktail of Cre-recombinase, Cre-silenced myr-tdTomato expression construct (control; magenta), Cre-activated myr-EGFP (*Bcl11a* null; cyan), and Cre-inducible Cre-recombinase. Two non-overlapping, genetically mosaic populations of control and *Bcl11a* null CPN_SL_ are generated. **(b)** Fluorescence images of P7 *Bcl11a*^flox/flox^ mouse brain electroporated *in utero* at E14.5 (control, magenta; *Bcl11a* null, cyan). High magnification greyscale images of contralateral cortex for control (b’) and *Bcl11a* null (b”). **(c)** Spatial distributions of ipsilateral transfection regions of mosaic CPN_SL_ populations at P7 show equivalent transfection coverage between control and *Bcl11a* null (control, n=3; *Bcl11a* null, n=3). **(d)** Quantification of contralateral axonal targeting at P7 reveals disruption of contralateral homotopic targeting of *Bcl11a* null CPN_SL_ axons compared to control (control, n=3; *Bcl11a* null, n=3). **(e)** Fluorescence images of P7 mouse brain reveal increased aberrant CPN_SL_ axonal projections through the AC upon *Bcl11a* deletion. High magnification greyscale images of AC for control (e’) and *Bcl11a* null (e”). Scale bars: 1 mm. All sections were immunolabeled for GFP (*Bcl11a* null; cyan) and RFP (control; magenta).

**Supplementary Figure 3:**
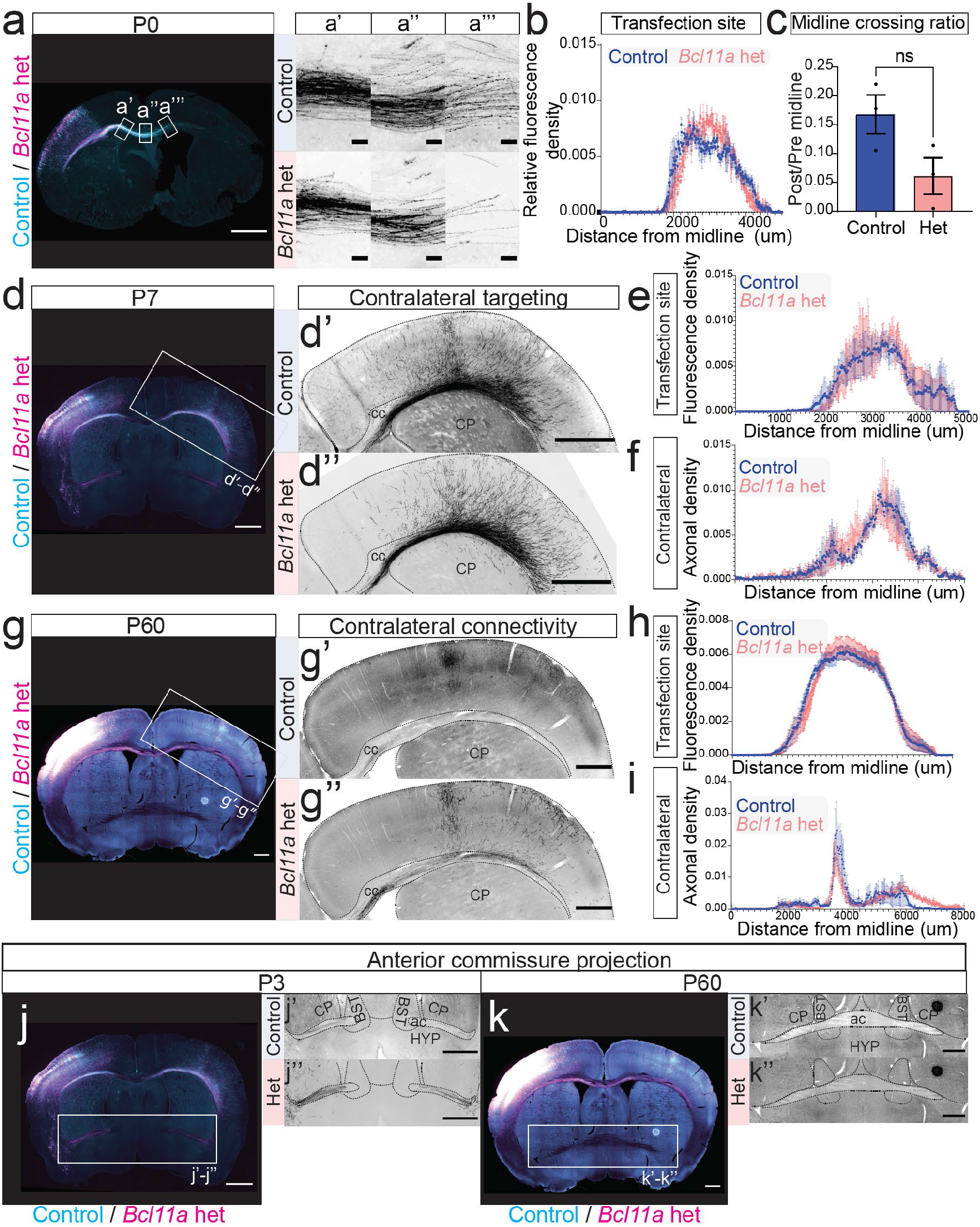
**(a)** Fluorescence images of P0 *Bcl11a*^flox/+^ mouse brain with mosaic populations of *Bcl11a* het (magenta) and control (cyan) reveals decreased number of *Bcl11a* het axons crossing the midline (a’’) and reaching the contralateral hemisphere (a’’’) compared to control, despite an equivalent number of pre-midline axonal fibers (a’). **(b)** Spatial distributions of ipsilateral transfection regions of mosaic CPN populations at P0 reveal equivalent transfection coverage between control and *Bcl11a* het. **(c)** Quantification of midline crossing ratio in somatosensory cortex reveals a trend for a lower percentage of *Bcl11a* het fibers crossing the midline compared to control (unpaired two-tailed Student’s t test; Control, n=3; *Bcl11a* het, n=3). **(d)** Fluorescence images of P7 *Bcl11a*^flox/+^ mouse brain, with mosaic populations of *Bcl11a* het (magenta) and control (cyan), electroporated *in utero* at E14.5. High magnification greyscale images of contralateral cortex for control (d’) and *Bcl11a* het (d”). Scale bars: 1 mm. **(e)** Spatial distributions of transfection regions at P7 reveal highly equivalent transfection coverage between control and *Bcl11a* het CPN somata (Control, n=3; *Bcl11a* het, n=3). **(f)** Quantification of contralateral axonal targeting at P7 reveals that contralateral homotopic targeting of *Bcl11a* het CPN axons remains unchanged compared to control (Control, n=4; *Bcl11a* het, n=4). **(g)** Fluorescence images of P60 mouse brain reveal similar axonal projections of *Bcl11a* het (g’) compared to control (g’’) in the contralateral cortex. Scale bars: 1 mm. **(h)** Spatial distributions of transfection regions at P60 reveal highly equivalent transfection coverage between control and *Bcl11a* het CPN somata (Control, n=6; *Bcl11a* het, n=6). **(i)** Quantification of contralateral axonal targeting at P60 reveals that contralateral homotopic targeting of *Bcl11a* het CPN axons remains mostly unchanged with slight increase in lateral projections compared to control (Control, n=6; *Bcl11a* het, n=6). **(j)** Fluorescence images of P3 mouse brain reveal increased aberrant CPN_SL_ axonal projections through AC upon *Bcl11a* heterozygous deletion. High magnification greyscale images of AC for control (j’) and *Bcl11a* het (j”). Scale bars: 1 mm. **(k)** Fluorescence images of P60 mouse brain reveal CPN_SL_ do not project through AC in adulthood in both control (k’) and *Bcl11a* het (k’’). Scale bars: 1 mm.

**Supplementary Figure 4:**
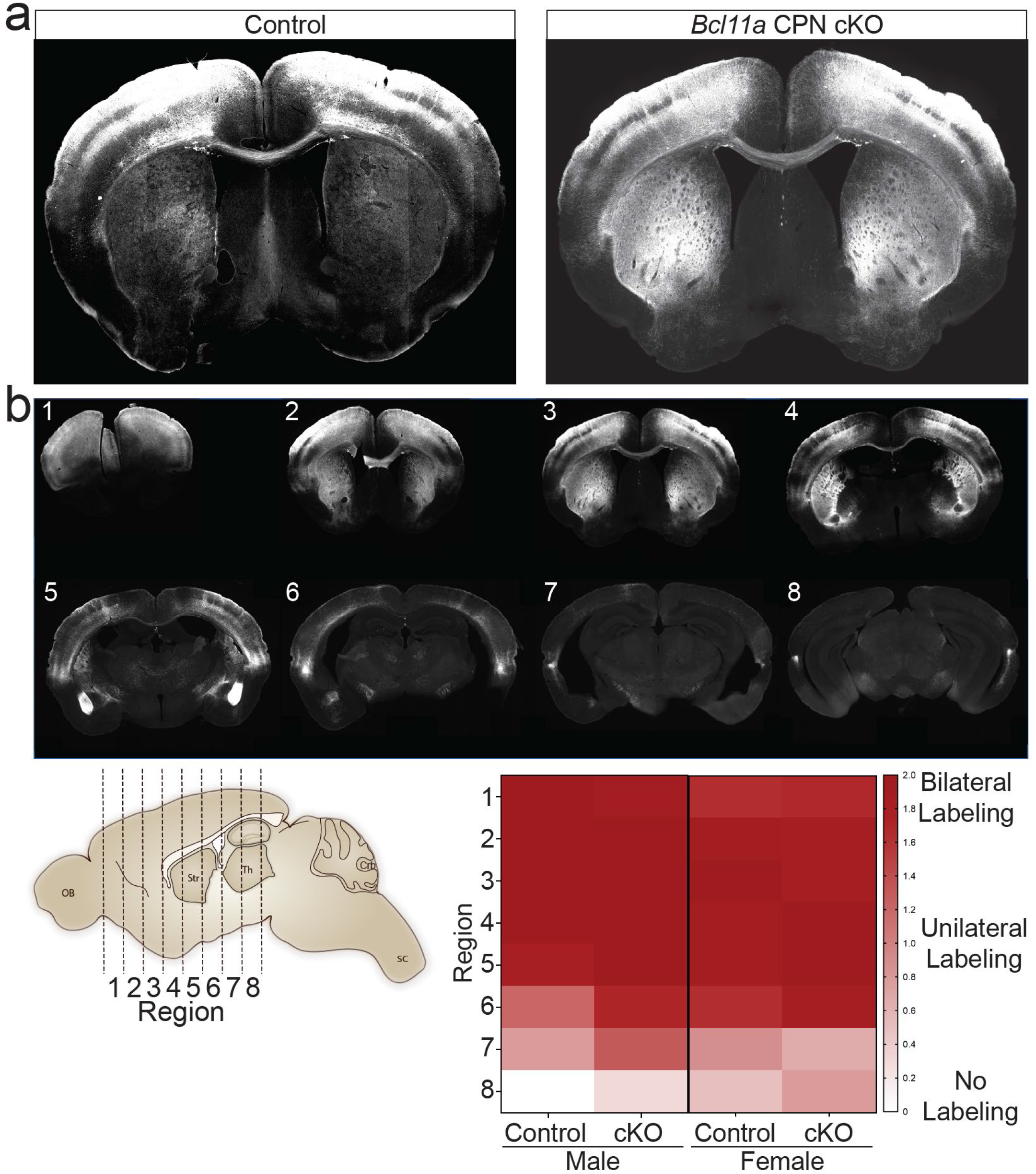
**(a)** Fluorescence images of P180 *Bcl11a*^flox/flox^ mouse brain bilaterally electroporated *in utero* at E14.5 with either myr-EGFP (control; left panel) or myr-tdTomato and Cre (*Bcl11a* CPN_SL_ cKO; right panel). **(b)** Successful bilateral targeting of CPN_SL_ was confirmed following the completion of behavioral experiments. Top: Eight rostral to caudal brain sections every ∼200 µm are shown from a representative mouse bilaterally electroporated at E14.5. Lower left: Schematic shows a sagittal view of the eight rostro-caudal section locations in each mouse. Lower right: Heatmap displays unitless fluorescence positivity. Each section is given either a score of “0” if both hemispheres don’t have any fluorescent signal, or a score of “1” if one of the hemispheres have a fluorescent signal, or a score of “2” if both hemispheres have fluorescent signal. The heatmap displays the average of scores for each section location per group (male control, male *Bcl11a* CPN_SL_ cKO, female control, and female *Bcl11a* CPN_SL_ cKO), revealing strong bilateral labeling rostro-caudally through sensorimotor cortex and section levels 1 through 5-6, and substantially reduced at visual cortex section levels 7-8.

**Supplementary Figure 5:**
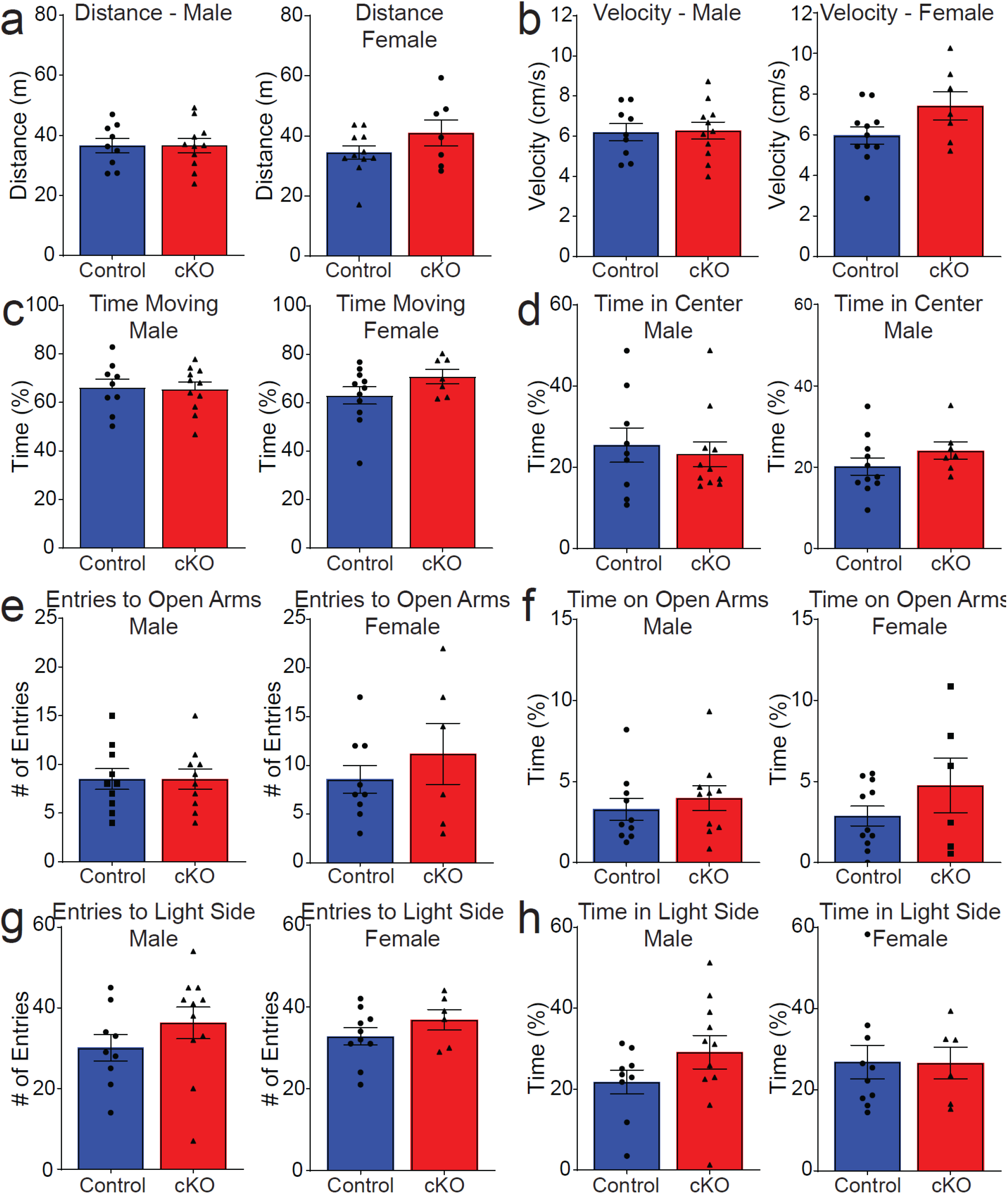
**(a-d)** Neither male nor female *Bcl11a* CPN_SL_ cKO mice displayed difference compared with control mice in distance moved (a), velocity (b), time in motion (c), or time spent in the center (d) in open field testing (two-tailed Student’s t test; control male, n=9; *Bcl11a* CPN_SL_ cKO male, n=11; control female, n=11; *Bcl11a* CPN_SL_ cKO female, n=7). **(e-f)** There is also no significant difference between *Bcl11a* CPN_SL_ cKO and control mice in the number of entries to open arms (e) and the time spent on open arms (f) in elevated plus maze test (two-tailed Student’s t test; control male, n=10; *Bcl11a* CPN_SL_ cKO male, n=10; control female, n=9; *Bcl11a* CPN_SL_ cKO female, n=6). **(g-h)** Similarly, both male and female *Bcl11a* CPN_SL_ cKO mice made a similar number of entries to the light chamber (g) and spent a similar amount of time in the light chamber (h) compared to control mice in light/dark chamber test (two-tailed Student’s t test; control male, n=9; *Bcl11a* CPN_SL_ cKO male, n=11; control female, n=10; *Bcl11a* CPN_SL_ cKO Female, n=6). ns, nonsignificant. n, number of distinct mice used. Results are represented as mean ± s.e.m.

**Supplementary Figure 6:**
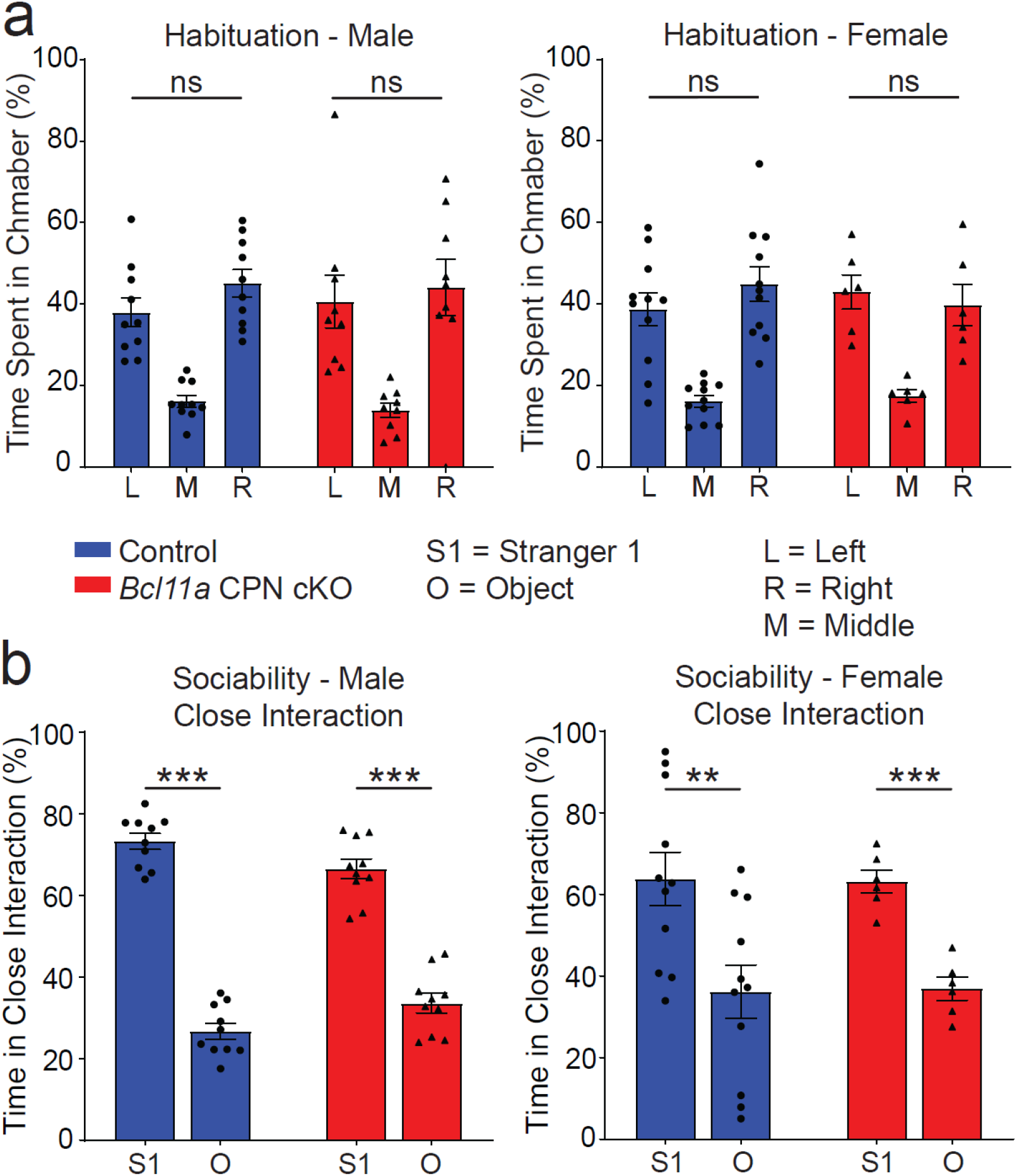
**(a)** Neither *Bcl11a* CPN_SL_ cKO nor control mice exhibited any preference for either of the side chambers during the 10-minute habituation period (neither males nor females; one-way ANOVA followed by Bonferroni multiple comparison test; control male, n=10; *Bcl11a* CPN_SL_ cKO male, n=8; control female, n=11; *Bcl11a* CPN_SL_ cKO female, n=6). **(b)** Both male (left) and female (right) *Bcl11a* CPN_SL_ cKO mice spent significantly more time in close interaction with the stranger mouse compared to the object (two-tailed Student’s t test; control male, n=10; *Bcl11a* CPN_SL_ cKO male, n=10; control female, n=11; *Bcl11a* CPN_SL_ cKO female, n=6). *, p-value<0.05; **, p-value<0.01; ***, p-value<0.001; ns, nonsignificant. n, number of distinct mice used. Results are represented as mean ± s.e.m.

**Supplementary Figure 7:**
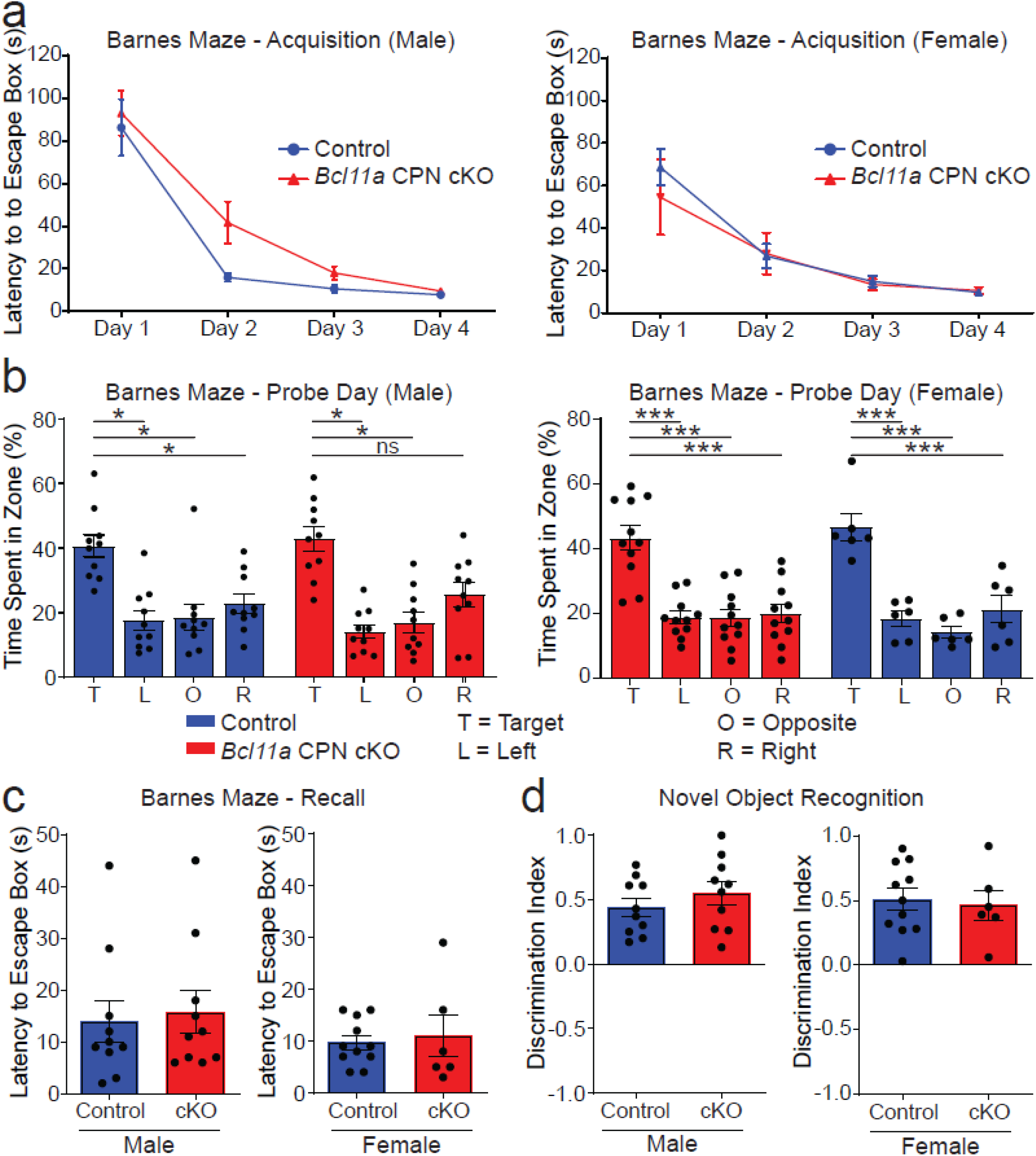
**(a)** *Bcl11a* CPN_SL_ cKO and control mice performed similarly in finding the escape box during the 4-day acquisition period of the Barnes maze test (both males and females; two-way ANOVA followed by Bonferroni multiple comparisons test; control male, n=10; *Bcl11a* CPN_SL_ cKO male, n=11; control female, n=11; *Bcl11a* CPN_SL_ cKO female, n=6). **(b)** Both *Bcl11a* CPN cKO and control mice spent significantly more time in the target quadrant on the probe day of the Barnes maze test (both males and females; one-way ANOVA followed by Bonferroni multiple comparisons test; control male, n=10; *Bcl11a* CPN_SL_ cKO male, n=10; control female, n=11; *Bcl11a* CPN_SL_ cKO female, n=6). **(c)** On the recall day of the Barnes maze test, *Bcl11a* CPN cKO and control mice spent similar amounts of time finding the escape box (both males and females; two-tailed Student’s t test; control male, n=10; *Bcl11a* CPN_SL_ cKO male, n=10; control female, n=11; *Bcl11a* CPN_SL_ cKO female, n=6). **(d)** Both male (left) and female (right) *Bcl11a* CPN_SL_ cKO mice spent more time interacting with the novel object compared to the familiar object (two-tailed Student’s t test; control male, n=10; *Bcl11a* CPN_SL_ cKO male, n=10; control female, n=11; *Bcl11a* CPN_SL_ cKO female, n=6). *, p-value<0.0332; **, p-value<0.0021; ***, p-value<0.0002; ns, nonsignificant. n, number of distinct mice used. Results are represented as mean ± s.e.m.

**Supplementary Figure 8:**
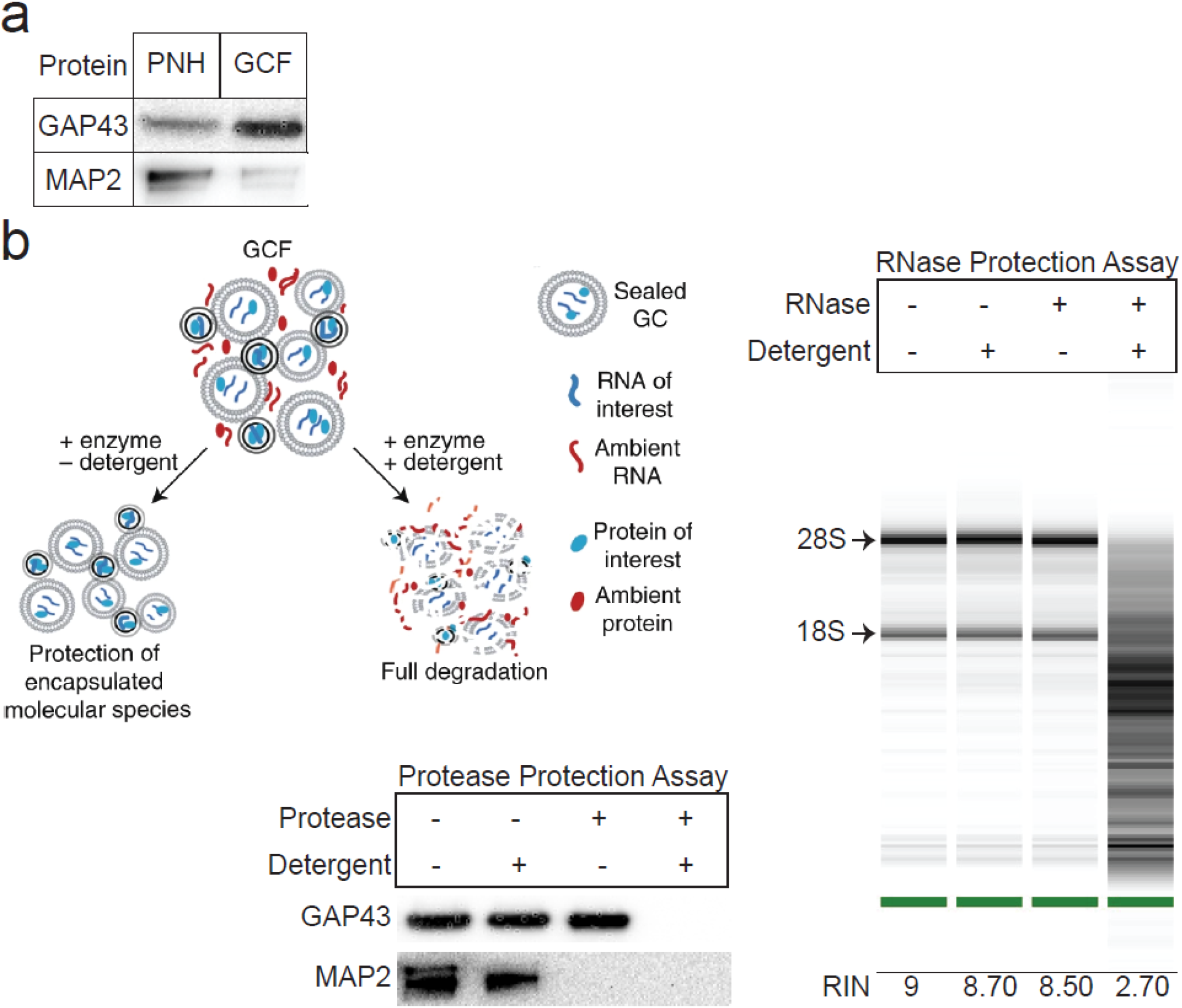
Subcellular fractionation enriches for intact, membrane-protected GCs. **(a)** Western blot of GAP43 and MAP2 proteins in PNH and GCF samples reveals substantial enrichment of the GC-marker GAP43 in the GCF sample and substantial de-enrichment of the somato-dendritic marker MAP2. **(b)** Schematic for, and results of, GC protection assays using Western blot for GAP43 and MAP2 proteins in GCF, and TapeStation of total RNA extracted from GCF. Neither detergent alone nor trypsin protease alone degrades GAP43, which is fully degraded with membrane permeabilization by detergent plus protease, indicating membrane protected contents, including the GC marker GAP43. In contrast, while detergent alone does not degrade the somato-dendritic MAP2, protease alone does, indicating that it is ambient and external to GCs, residual from the tissue homogenizing process designed to leave GCs intact. Similarly, neither detergent nor RNase alone degrade the GC-enclosed RNA, while the combination does, reinforcing that the resultant GCs are membrane protected, and contain non-degraded RNA.

**Supplementary Figure 9:**
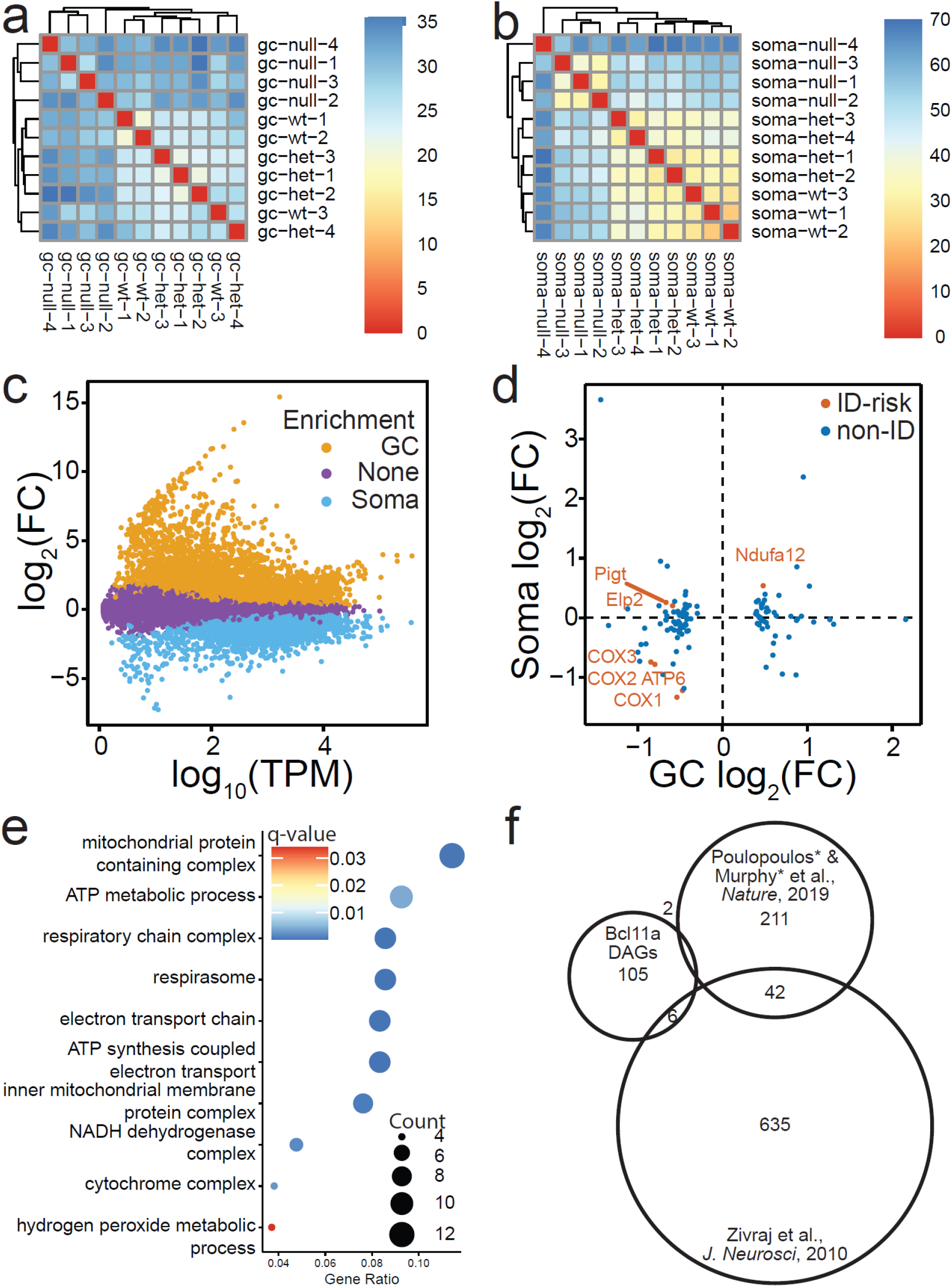
Differential enrichment analysis is supported by high-quality RNA sequencing, and reveals differentially abundant genes (DAGs) across *Bcl11a* genotypes and CPN subcellular compartments. **(a)** Clustering of GC samples, revealing expected, quality clustering of like sample. Lower sample-to-sample distances have warmer colors, and higher sample-to-sample distances have colder colors. **(b)** Clustering of soma samples, again revealing expected, quality clustering of like samples. Lower sample-to-sample distances have warmer colors, and higher sample-to-sample distances have cooler colors. **(c)** MA-plot displaying changes in gene abundance in CPN_SL_ GCs relative to CPN_SL_ somata. GC-enriched genes are orange, soma-enriched genes are blue, and unenriched genes are purple. **(d)** Quadrant plot comparing changes in abundance of GC DAGs in CPN_SL_ somata and CPN_SL_ GCs after deletion of *Bcl11a*. DAGs are colored by their ID classification – ID risk genes are red, and non-ID risk genes are blue. **(e)** GO enrichment analysis of DAGs. Points are colored by q-value and sized by number of genes per term. **(f)** Venn diagram comparing GC DAGs identified in this study to core axon GC molecules described by our lab (for CPN; Poulopoulos*, Murphy* et al., *Nature*, 2019) and the C. Holt lab (for mouse and xenopus retinal ganglion cells; Zivraj et al., *J. Neurosci*, 2010).

**Supplementary Figure 10:**
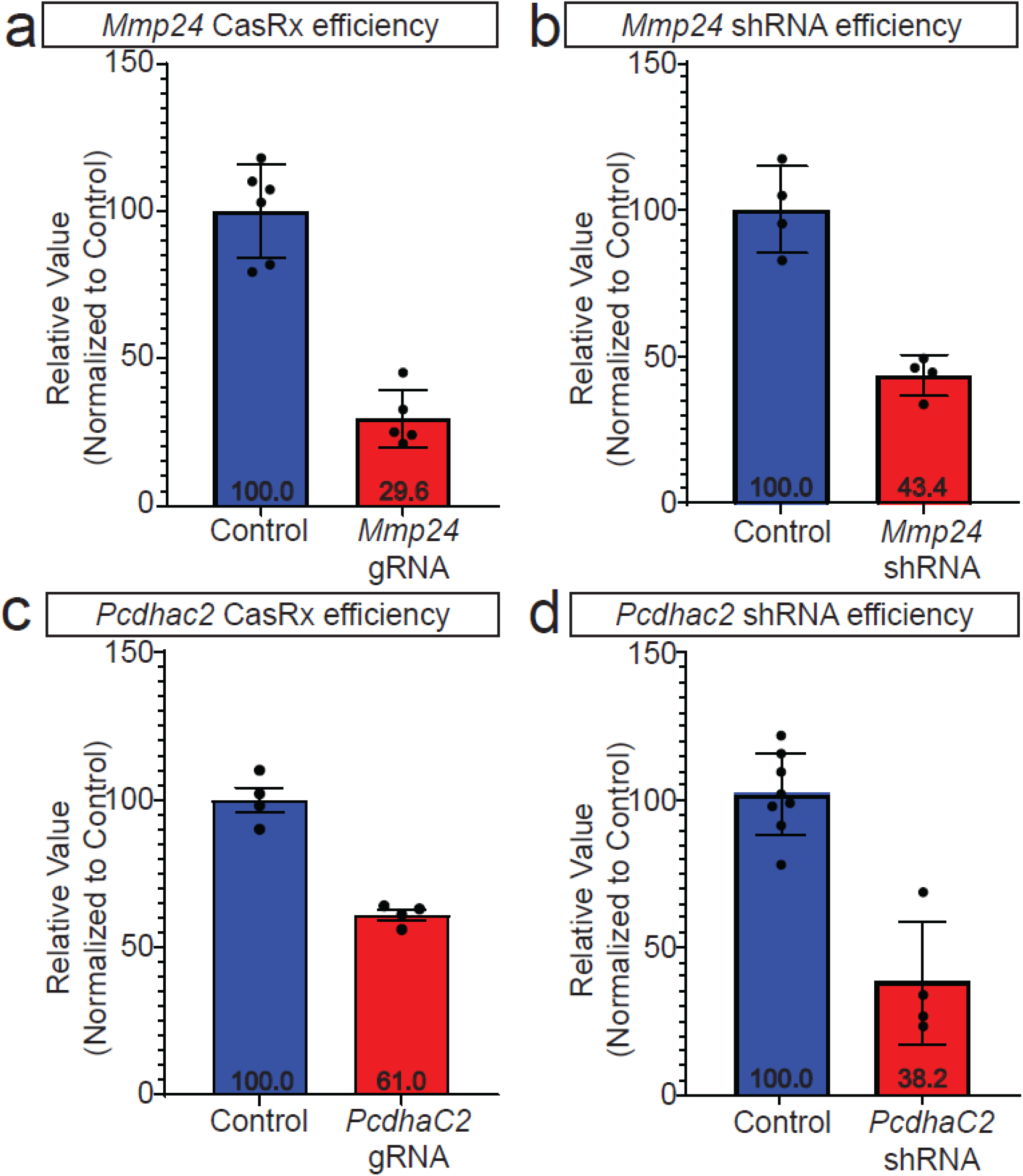
**(a)** *Mmp24* expression by CPN_SL_ is efficiently reduced using the CRISPR-CasRx system, assessed by qRT-PCR (control, n=6; *Mmp24 gRNA*, n=5). **(b)** *Mmp24* expression by CPN_SL_ is moderately efficiently reduced using the *Mmp24* shRNA assessed, by qRT-PCR (control, n=4; *Mmp24 shRNA*, n=4). **(c)** *PcdhαC2* expression by CPN_SL_ is reduced using the CRISPR-CasRx system, assessed by qRT-PCR (control, n=4; *PcdhαC2 gRNA*, n=4). **(d)** *PcdhαC2 shRNA* efficiently reduces *PcdhαC2* expression by CPN_SL_, assessed by qRT-PCR (control, n=8; *PcdhαC2 shRNA*, n=4). For all experiments, i*n utero* electroporation was performed at E14.5 to target CPN_SL_, and labeled CPN_SL_ were sorted at postnatal day 3 (P3). “n” refers to number of the distinct sorted P3 CPN_SL_ samples.

**Supplementary Figure 11:**
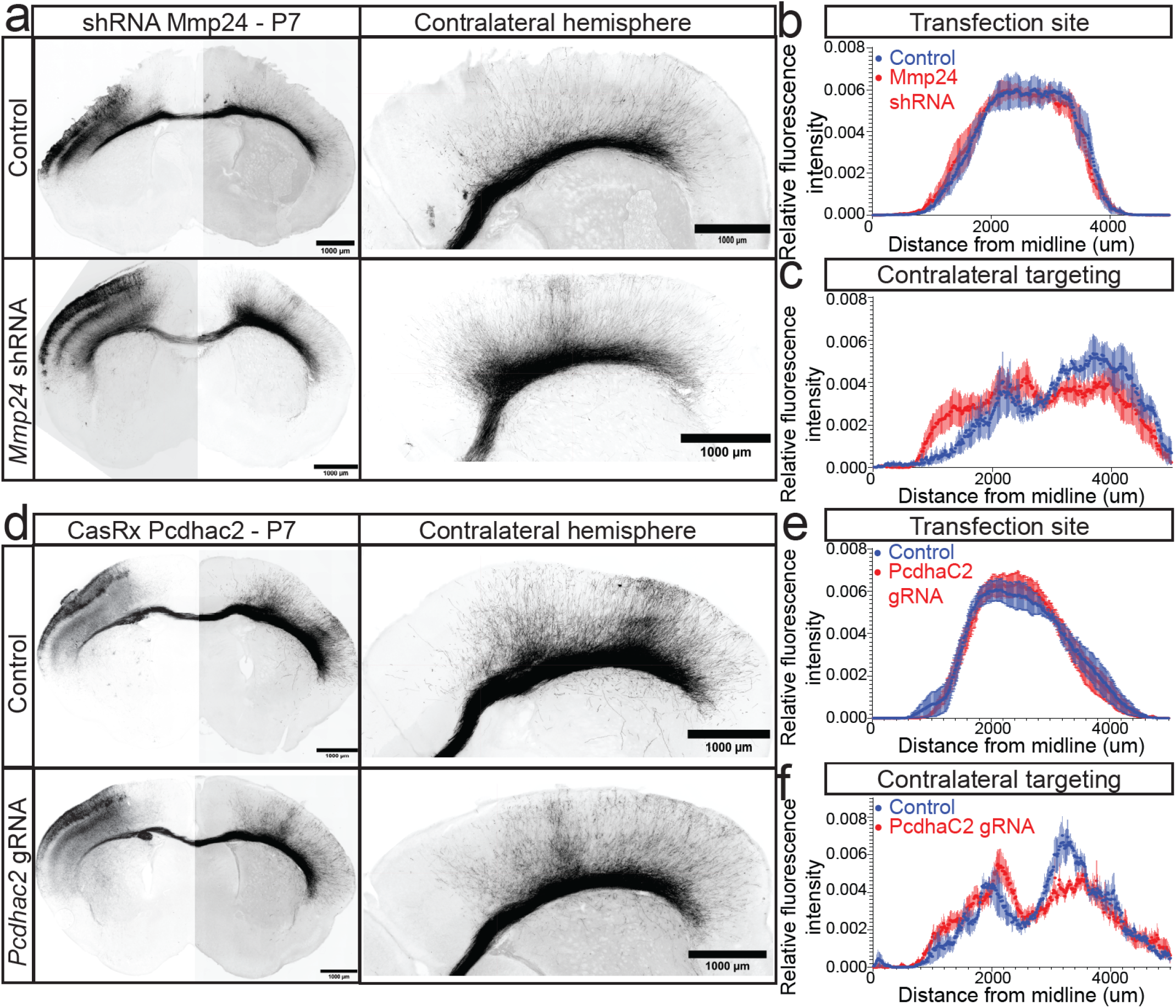
**(a)** Greyscale images of P7 wildtype mouse brains electroporated *in utero* at E14.5 with either a non-targeting (control) or *Mmp24*-directed shRNA (*Mmp24* shRNA), and a myr-tdTomato expression construct. Right panels show high magnification images of contralateral cortex for control (upper panel) and *Mmp24* shRNA (lower panel). Scale bars: 1 mm. **(b)** Spatial distributions of ipsilateral targeting reveal highly equivalent mediolateral transfection coverage between control and *Mmp24* shRNA CPN_SL_ somata via *in utero* electroporation (control, n=5; *Mmp24* shRNA, n=6). **(c)** Quantification of contralateral axonal targeting at P7 reveals disruption of homotopic targeting of *Mmp24* shRNA CPN_SL_ axons compared to control (control, n=5; *Mmp24* shRNA, n=5). **(d)** Greyscale images of P7 wildtype mouse brains electroporated *in utero* at E14.5 with either a non-targeting (control) or *PcdhαC2*-directed guide RNA (*PcdhαC2* gRNA), and a myr-tdTomato expression construct. Right panels show high magnification images of contralateral cortex for control (upper panel) and *PcdhαC2* gRNA (lower panel). Scale bars: 1 mm. **(e)** Spatial distributions of ipsilateral targeting reveal highly equivalent mediolateral transfection coverage between control and *PcdhαC2* shRNA CPN_SL_ somata via *in utero* electroporation (control, n=6; *PcdhαC2* gRNA, n=9). **(f)** Quantification of contralateral axonal targeting at P7 reveals disruption of homotopic axonal targeting of *PcdhαC2* gRNA CPN_SL_ axons compared to control (control, n=6; *PcdhαC2* gRNA, n=7). “n” refers to the number of distinct mice used. All sections were immunolabeled for RFP.

